# Reliable data collection in participatory trials to assess digital healthcare apps

**DOI:** 10.1101/2020.02.05.935049

**Authors:** Junseok Park, Seongkuk Park, Kwangmin Kim, Gwangmin Kim, Jaegyun Jung, Sungyong Yoo, Gwan-su Yi, Doheon Lee

## Abstract

The number of digital healthcare mobile apps on the market is increasing exponentially owing to the development of the mobile network and widespread usage of smartphones. However, only a few of these apps have undergone adequate validation. As with many mobile apps, healthcare apps are generally considered safe to use, making them easy for developers and end-users to exchange them in the marketplace. The existing platforms are not suitable to collect reliable data for evaluating the effectiveness of the apps. Moreover, these platforms only reflect the perspectives of developers and experts, not of end-users. For instance, data collection methods typical of clinical trials are not appropriate for participant-driven assessment of healthcare apps because of their complexity and high cost. Thus, we identified a need for a participant-driven data collection platform for end-users that is interpretable, systematic, and sustainable —as a first step to validate the effectiveness of the apps. To collect reliable data in the participatory trial format, we defined distinct stages for data preparation, storage, and sharing. Interpretable data preparation consists of a protocol database system and semantic feature retrieval method to create a protocol without professional knowledge. Collected data reliability weight calculation belongs to the systematic data storage stage. For sustainable data collection, we integrated the weight method and the future reward distribution function. We validated the methods through statistical tests conducted on 718 human participants. The validation results demonstrate that the methods have significant differences in the comparative experiment and prove that the choice of the right method is essential for reliable data collection. Furthermore, we created a web-based system for our pilot platform to collect reliable data in an integrated pipeline. We validate the platform features with existing clinical and pragmatic trial data collection platforms. In conclusion, we show that the method and platform support reliable data collection, forging a path to effectiveness validation of digital healthcare apps.

## 1 Introduction

Digital healthcare has become one of the most promising fields in the healthcare industry with the widespread popularity of wireless devices such as smartphones. Notably, approximately 200 mobile health apps are newly added on a daily basis in mobile app stores, and investments in digital health are booming [22, 80]. While many of these apps are general wellness applications that help users to manage their activities, some healthcare apps claim to improve a symptom or disease directly. The treatments proposed by the apps take the form of information or visual stimuli, which are effectively delivered to users as a digital therapy. reSET-O, an 84-day prescription digital therapeutic app for the treatment of opioid use disorder, has been approved by the U.S. Food and Drug Administration (FDA) [46]. Akili Interactive has clinically demonstrated that interactive digital treatment through a video game, which is under review by the FDA, may improve symptoms of attention deficit hyperactivity disorder (ADHD) and sensory processing disorder [2, 106]. Omada Health showed that a web-based diabetes prevention program could significantly lower hemoglobin A1c levels. Moreover, the web-based program showed a higher rate of patient engagement than the traditional in-person diabetes prevention plan [84]. Their product has received tremendous attention from researchers and the public as an example of digital medicine [100].

The increased interest in digital health has led researchers to study the effectiveness of current apps. As a practical example, Pokémon GO, which is a game app based on augmented reality, has positive effects on social interaction: 43.2% of people spent more time with their family [52]. In contrast, a 6-week online intervention experiment has revealed that Headspace, which is a healthcare app for mindfulness by guided meditation, has relatively small effects on mindfulness [71]. The outcome contradicts the results of previous randomized controlled trials for the app [44]. Because of such contradictions, there is a need for a reliable mechanism to identify validated digital health apps [62].

Currently, there is no proper platform that evaluates the effectiveness of an app using a systematic and objective validation method in digital health [62]. In particular, mobile health apps have a very fast development cycle and are low-safety issue technology, which tends to reduce the burden of regulation. As a result, these apps have a direct trade characteristics between developers and end-users. However, the few existing platforms are not suitable to evaluate the effectiveness of the apps in this case. For example, review platforms and guidelines are emerging to solve the problem, but they only provide limited effectiveness information obtained from the perspectives of developers, experts, and regulatory authorities [19, 38, 56, 69]. Thus, we identified a need for a participant-driven data collection platform for end-users as an interpretable, systematic, and sustainable tool to validate the effectiveness of the apps.

An appropriate validation method begins with the collection of reliable data; next, data analysis is performed for verification [6]. Hence, data collection is the basis of data analysis. The utilization conditions of healthcare apps are different from an ideal research-intensive environment, and the collected data relate to the term “real-world data” [88]. An incomplete collection of real-word data can cause inaccurate data analysis due to inconsistency, poor data quality, and noisy or missing data [108]. Thus, the data collection is an essential stage of the verification workflow. We compare in the following previous expert-oriented data collection platforms in existing research fields to develop a participant-driven data collection platform.

A general study with high effectiveness and efficacy of the measurement method is a clinical trial. A clinical trial includes a reliable data collection procedure based on a clinical protocol in a controlled environment [79]. The clinical protocol is a translational guide for data collection, and advanced digital technology has been developed as an electronic data capture (EDC) platform to prepare, store, share, examine, and analyze data from electronic case report forms (eCRF) within the protocol. Therefore, the platform is complex for the public to use. For instance, Rave, EDC platform made by Medidata, showed the highest level of popularity and satisfaction in the G2 Crowd, a software comparison website [70]. Rave follows a typical EDC design approach. Besides, Rave has a more advanced feature to design the basic document structure of a protocol and provides optimized protocols using a PICAS database with trial cost information [34]. Despite this functionality, Rave is still appropriate for expert use. Transparency Life Sciences (TLS) is a leader in digital drug development services offering virtual and hybrid clinical trials [57]. TLS uses the crowdsourcing method for protocol design. In detail, experts create a survey project for a disease and release the project to all TLS users. After the release, the users interested in the project start to participate with their identities such as patient family members, researchers and patients. At the last step of the project, the TLS clinical trial team designs the protocol from the collected data of the participants. However, also in this case the creation of the protocol is driven by an expert team, rather than common users. Accordingly, the methods for data collection in clinical trials are not appropriate for participant-driven assessment of healthcare apps (which are exponentially growing) because of their complexity and high cost [24, 94].

A study on the real-world measure of the effectiveness of intervention in broad groups is a pragmatic trial [28]. The data collection platform of a pragmatic trial includes not only EDC but also specialized research tools and general surveys. These data collection platforms can be web-based survey applications, or mailed questionnaires, or specific healthcare apps developed from research kits [8, 12, 15]. Therefore, while the platforms still have the issue of complexity, there is also the possibility of collecting less reliable data. For instance, PatientsLikeMe is a platform that shares experience data for participants to understand possible treatments of particular disease conditions based on the experiences of others [101]. However, PatientsLikeMe does not provide an environment for the public to lead the study preparation, and the platform has no feature to evaluate the reliability of collected data. Another example is Amazon Mechanical Turk (MTurk). MTurk is a crowdsourcing marketplace for the recruitment of research participants and a platform for conducting surveys [73]. However, the platform does not provide any standardized method at the data preparation stage. In other words, the platforms need clinical experts to prepare data collection procedures and system based on their knowledge. MTurk provides a feature to approve or reject an individual worker, but the feature relies on a subjective judgement with obvious objectivity limitations. We found that this platform has no suitable method to measure the reliability of the data collected from the participants.

The participatory trial platform allows a public user to create and conduct the participant-driven study to measure the effectiveness of products within daily life. The core factors of the platform are simplicity, reliability and sustainability. Based on the definition, we identified comparable platforms that have an alternative, similar or essential feature. For example, Google Play and Apple App Store are alternative platforms because they keep user ratings and reviews in the public domain [3, 35, 66]. Both platforms have free text app reviews and scaled rating functions as a data storage feature. However, the review feature has a natural language processing problem, which is not structural data collection [61]. In other words, the platforms have no simple data preparation method to create a systematic data collection protocol. In addition, the platforms have a possible risk of transfer biases that could affect new reviews and ratings, because they expose previously collected data to new participants [72]. Another limitation of the platforms is that they do not offer any features to evaluate data reliability. RankedHealth and NHS Apps Library can also represent similar platforms [1, 36, 90]. RankedHealth has a procedure to minimize the influence of individual reviewers for data reliability. NHS App Library publishes safe and secure apps through questions designed by experts from technical and policy backgrounds. However, these platforms have a limitation. The consensus of the experts is focused on evaluating the technical performance of an app. Expert assessment does not validate the effectiveness of user experience of the app. Thus, they are not appropriate for participant-driven studies.

Finally, all the platforms we mentioned have limitations that do not reflect the prevention of drop-out and software characteristics of digital healthcare apps. A study with a daily collection of pain data using digital measures has shown that the self-reporting completion rate is at an average of 71.5% (261/365) days. The latest researches are attempting to develop an incentivized program or in-game rewards to increase the self-reporting rates because sustainable data sharing for collecting large amounts of data is a crucial function of the participatory trial platforms [43, 49, 92]. Furthermore, unlike drugs, functional foods, and mechanical devices that are difficult to modify after the market launch, healthcare apps can be potentially updated as software [26]. The app features may require iterative evaluation following the upgrade. In summary, a new platform for digital healthcare apps should have a sustainable data collection function.

Here, we propose a participant-driven reliable data collection method for participatory trial platforms as a first stage to understand the effectiveness of healthcare apps. The method consists of three steps: understandable data preparation, systematic data storage and sustainable data sharing. We utilize a clinical trial protocol database and a semantic relatedness method for the participatory trial protocol to prepare data collection. We develop a data weight reliability formula that collects data systematically. We propose a future reward distribution function connected to data reliability weight for sustainable collection of data. In the results section, we describe the following experiments to validate the reliable data collection method: comparison to the simplicity of data preparation, data reliability weight validation and future reward distribution effect observation. We report testing on a total of 718 human participants for the experiments. The Institutional Review Board (IRB) of KAIST approved an IRB exemption for the experiments.

Moreover, we have developed a web-based pilot platform accessible to the public with real-world data as a crowdsourcing tool based on the citizen science concept. The pilot platform systematically integrates all proposed methods. We conduct case studies on the pilot platform to validate efficient recruitment. To demonstrate the advantages of the proposed platform, we compare the platform functionality to existing platforms.

## 2 Definition of Participatory Trial

A participatory trial is an expanded form of a human-involved trial in which the public is used to test the effectiveness of products within their daily life. The concept follows crowdsourcing and citizen science in the aspect of data-driven science [40, 53]. The participatory trial includes voluntary participation, unlike selective inclusion of the clinical trial and broad inclusion of the pragmatic trial. The participants from the public operate the trials. The objective of the participatory trial is to inform consumers about the effectiveness of a product, and the method consists of a protocol reflecting daily life care to maximize the reliability of the clinically relevant results. We have compared the continuum between clinical trials, pragmatic trials, and participatory trials in Table 1 [67]. We have defined a participatory trial to ameliorate current trial definitions in modern society.

**Table 1.**
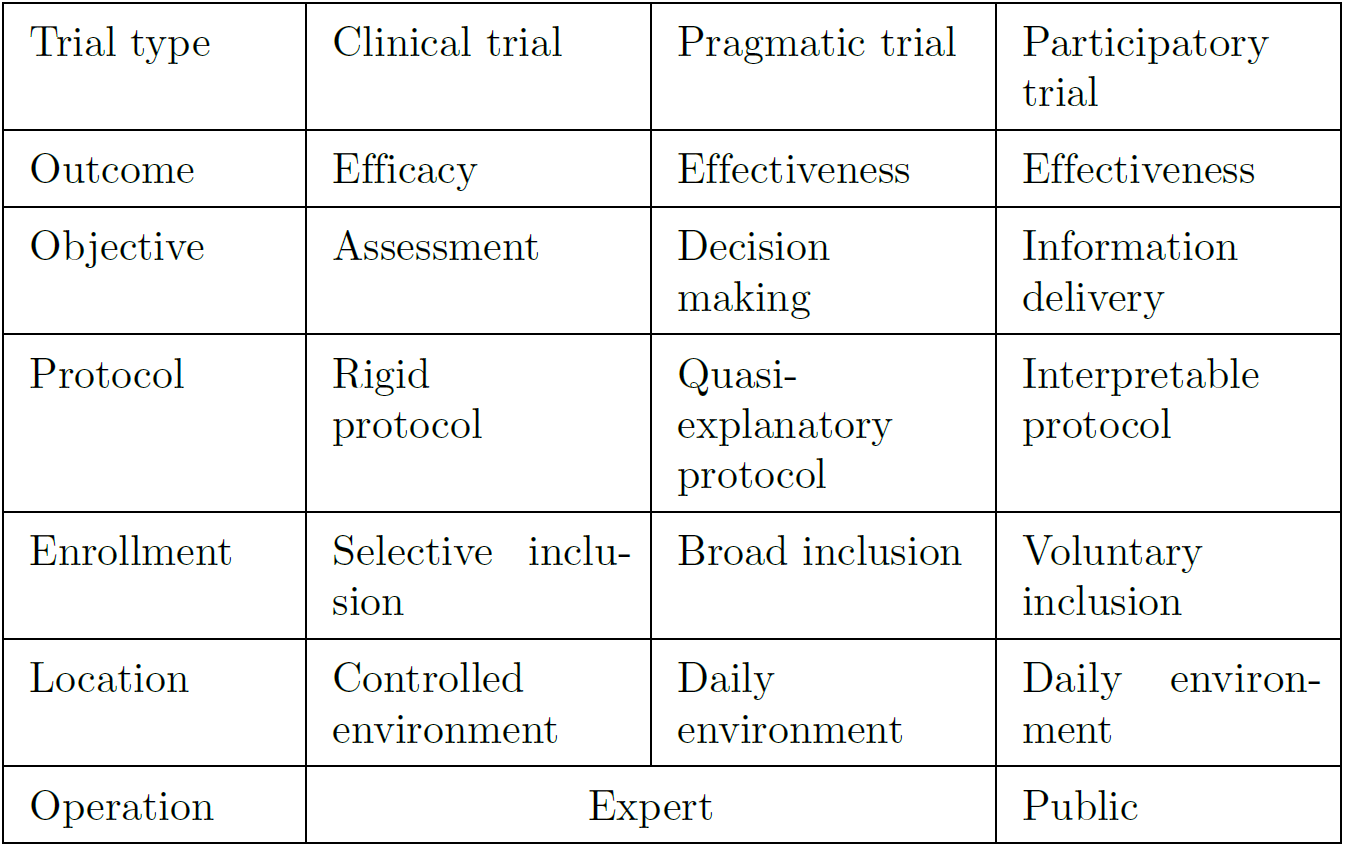
Comparison between current trials and participatory trial

Limited to this manuscript, we also defined the meaning of a word or phrase as following;

- Platform: Digital platform, that is an environmental software system associated with each functional part of application programs.
- Data preparation: Determining the data collection structure and having the same meaning as designing the protocol.
- Data storage: Evaluating collected data to store reliably and to determine the value of the data.
- Data sharing: Integrating collected data to share with others and to provide benefit with the data provider.

## 3 Related Work

### 3.1 Clinical Trials and Platforms

Clinical trials have been the gold standard for evaluation or development of medical interventions for over 70 years [9]. Focused on evaluating the efficacy of specific interventions, clinical trials often use controlled settings and strict criteria for participants and practitioners. The EDC system is increasingly recognized as a suitable method that has advantages in real-time data analysis, management, and privacy protection [99]. The system is most useful in trials with complicated contexts, such as international, multi-centered, and cluster-randomized settings [96]. However, because of their nature of strictness and complexity, clinical trials still have certain limitations in the thorough evaluation of effectiveness or external validity [81].

### 3.2 Pragmatic Trials and Platforms

Clinical trials are often time-consuming and challenging because they utilize rigorous methods. These characteristics of clinical trials have increased the need for pragmatic trials that seek to understand real-world evidence, assessing the effectiveness of the intervention in actual clinical practice settings [28]. As introduced in the “PRECIS’’ assessment by Thorpe et al., these trials try to maximize the external validity of the research by including the heterogeneous population and setting patient-centered endpoints [93]. Various platforms, including mailed questionnaires, web-based forms, and research kits, are used for these kinds of trials [8, 12, 15]. Consequently, pragmatism is an emerging source of clinical evidence, from pediatric asthma and cardiovascular diseases to monetary incentives for smoking cessation [4, 36, 85]. However, it is subject to several challenges such as patient recruitment, insufficient data collection, and treatment variability [28, 91]. In addition, inconsistency in data platforms is another major limitation of pragmatic trials [95].

## 4 method

We divided the data for evaluating the effectiveness of the healthcare app into the stages of interpretable preparation, systematic storage, and sustainable sharing. We used the essential methods of each stage to collect highly reliable data continuously. In addition, from a participatory trial perspective, we have organized each method so that the public could easily participate and organize their own research, provide reliable information, and share it to induce continued interest (Figure 1).

**Figure 1.**
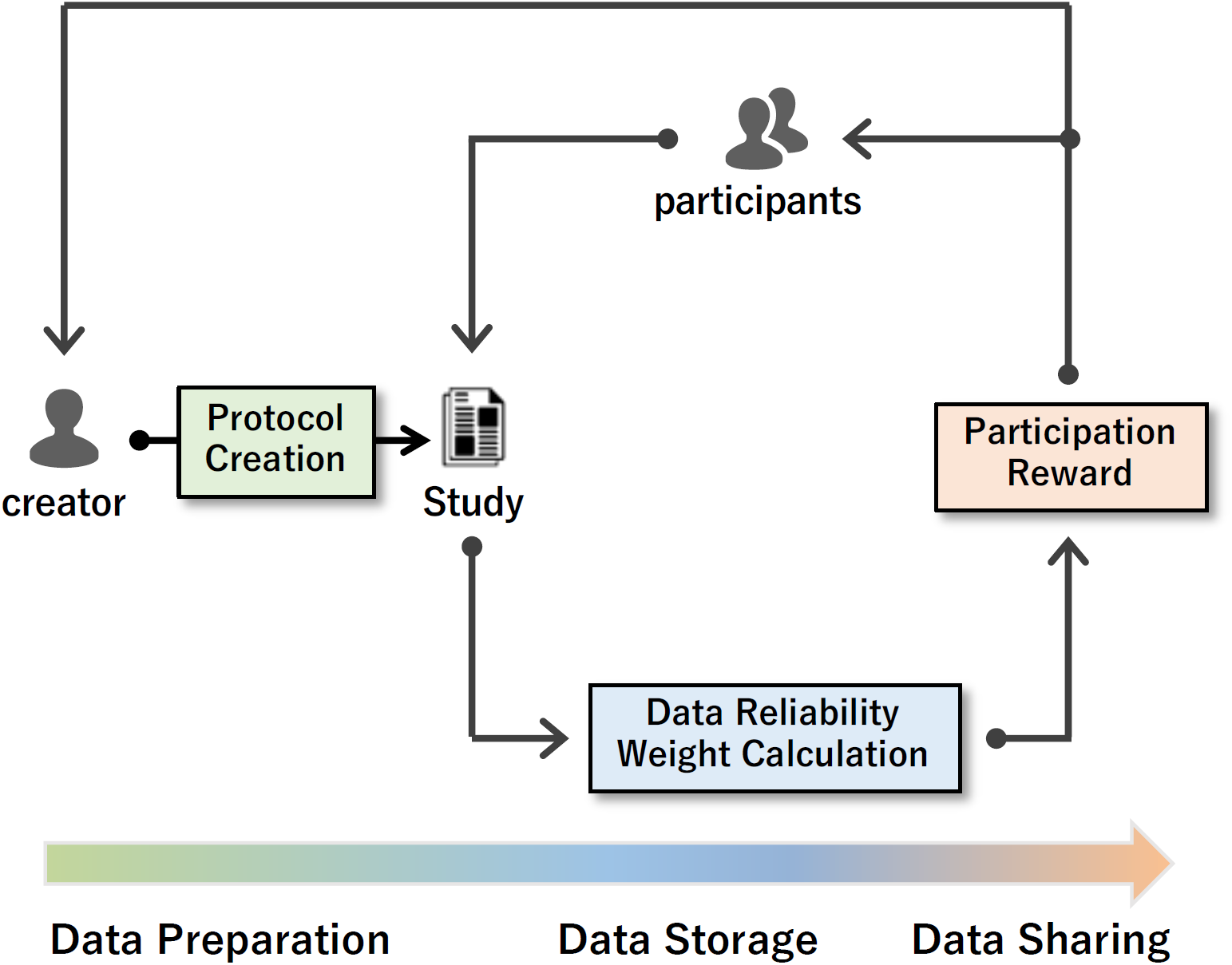
Method overview. The method consists of three parts: data preparation, data collection, and data distribution. In data preparation, a creator can devise a new study for the evaluation of an app by searching from the protocol creation method. Next, in data storage, the participants conduct a study, and the platform stores all the participants’ replies. At the same time, it collects statistics of the replies and calculates the data reliability weight of each reply. After finishing the study, in data sharing, the platform calculates a future reward depending on each person’s reliability of both the participants and the creator and distributes the reward to each of them.

### 4.1 Interpretable Data Preparing

At the data preparation stage, we determined what kinds of data should be collected, how to collect the data, and from where we would get the data. We had to consider the data as an interpretable resource to determine the effectiveness of a healthcare application. In the fields of clinical and pragmatic trials, researchers usually develop a protocol to facilitate the assessment of a changing symptom or disease [16, 93]. Healthcare applications also focus on improving a symptom or treating a disease. A phenotype is an observable physical characteristic of an organism, including a symptom and disease. Therefore, we can effectively measure the change of a phenotype from an existing protocol that is semantically related to the phenotype.

We developed a protocol database system and a concept embedding model to calculate semantic relatedness for a protocol retrieval system for interpretable data preparation. The database contains 184,634 clinical trial protocols for context-dependent protocol-element-selection [74]. We created the model to find a vector on a latent space based on distributed representations of a documented method. Furthermore, we calculated the semantic relatedness between input phenotype and protocols with cosine-similarity [55, 75]. Finally, we integrated the methods into the proposed platform for the participatory trial, so that anyone can create a study to easily determine the effectiveness of a healthcare application. Figure 2 shows the overall procedure of the method.

**Figure 2.**
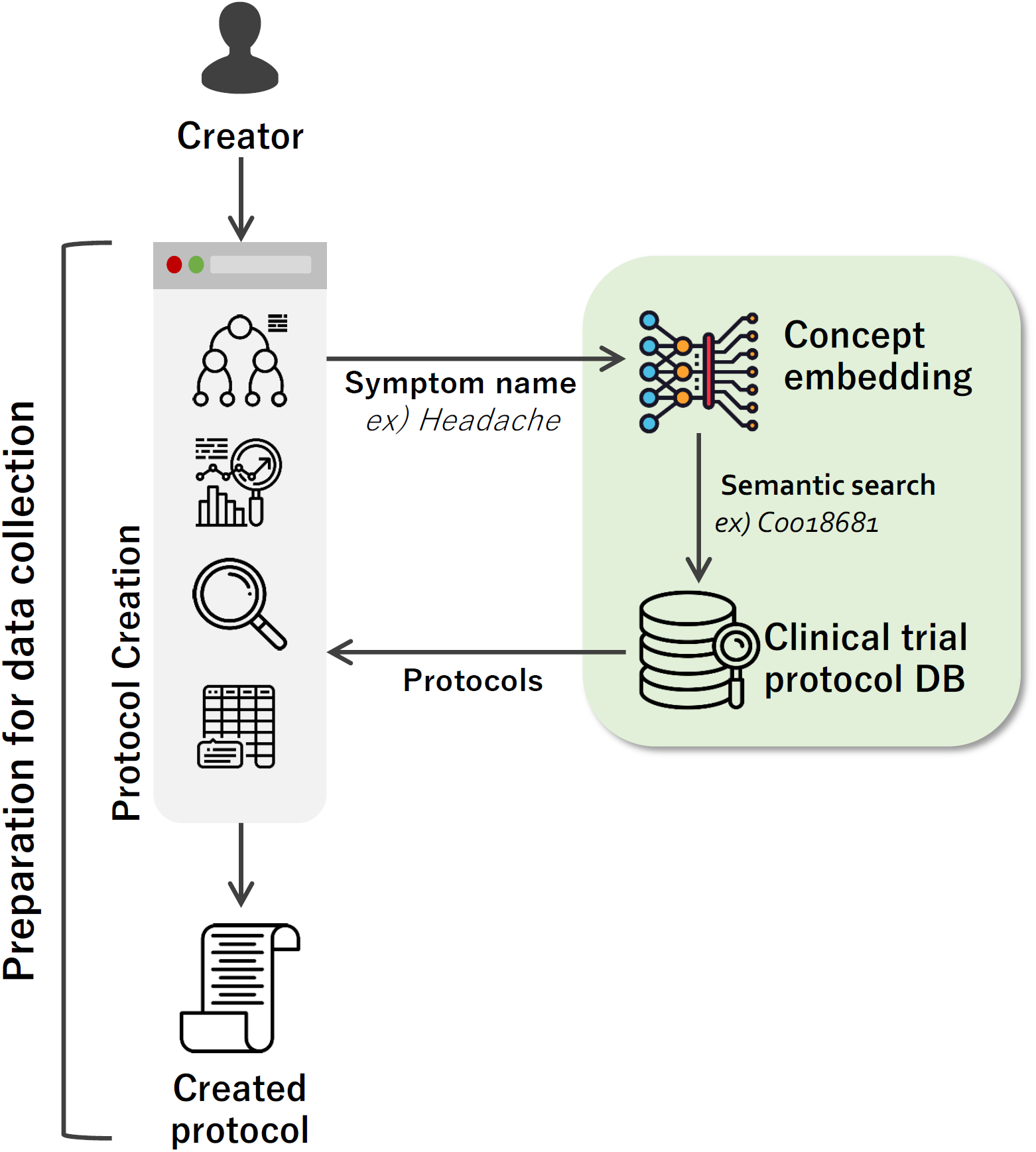
Data preparation to collect. The creator retrieves the previous protocol based on the symptom name of the app to verify effectiveness. Then the creator prepares to collect data on the retrieved protocol and creates the participatory trial protocol for data storing.

### 4.2 Systematic Data Storage

The participatory trial platform uses crowdsourcing methods to collect data. Crowdsourcing refers to obtaining data and using it for scientific research through voluntary participation from the public. Data collection based on crowdsourcing is applied in many research fields because it is a method that can quickly obtain a large amount of data from the public at a low cost [48, 51]. However, data collected from crowdsourcing is often pointed out to be less reliable than results obtained by systematic approaches [25, 59]. Therefore, devising a method to measure data credibility is the main challenge of the data reliability issue.

The primary purpose of this method is reliable data collection and delivery for effectiveness evaluation, which is the next level on the path to validation. To achieve this purpose, we needed a method to guarantee the reliability of input data. Therefore, we proposed a formula to calculate the data reliability weight (DRW) through combined effort measures of input data.

Careful, relevant, and purposive data input by the participant can improve the quality of data in systematic data storage [63]. Huang et al. developed an insufficient effort responding (IER) detection method to determine the input data needed for providing accurate responses and correctly interpreting the data [42]. The recommended composition of IER is response time, long-string analysis, and individual reliability. Response time (RT) is a method to determine the effort based on execution time [104]. Long-string analysis (LSA) uses intentional long-string patterns to assess the effort of a participant [21]. Individual reliability (IR) divides a pair of data items into two halves and uses the correlation between two halves to determine the normal response [83]. In addition, Huang et al. also presented a single self-report (SR) IER item, which includes empirical usefulness [41]. The SR is a single item for detecting IER, and SR consists of the same 7-point Likert scale [65]. We developed the DRW calculation method based on the IER methods.

We calculated *DRW*_*idx*_(*X*) according to the type of electronic case report form (eCRF) for obtaining DRW (1). *f*_*RT*_, *f*_*LSA*_, *f*_*SR*_ and *f*_*IR*_ are the indexes of DRW. The created protocol contains eCRFs, and an eCRF is a tool used to collect data from each participant. We provided four types of eCRF as described in Supplementary Table 1. We calculated DRW from the symptom and data type eCRF. The symptom type measures the presence and the degree of targeted symptoms only once. The data type measures the effectiveness of the targeted apps repeatedly. *X* = {*x*_1_, *x*_2_, …, *x*_*n*_} is the set of items in eCRF. The platform receives 11 types of item data in eCRF, as shown in Supplementary Table 2. Type codes ED, TEL, SET, DATE, PDATE, and FILE were not used to calculate the DRW.

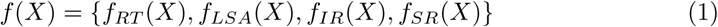

We calculated *f*_*RT*_ and *f*_*LSA*_ for symptom and data type of *X. f*_*RT*_ calculates the difference between the start and end time when the participant input the data to *X. f*_*LSA*_ uses the length input strings. On the other hand, we calculated *f*_*SR*_ and *f*_*IR*_ only for the symptom type of *X*. We did not include the SR and IR item in the data type of *X* to avoid distortion of the response by the participant on the iterative procedures. To measure whether a participant is attentive while entering data at the end, we placed an item of SR at the end of the symptom type of *X*. The measured pattern of participants should be similar to the symptom type. To reflect the pattern for *f*_*IR*_, we calculated a Pearson correlation coefficient [30]. *r*_*IR*_ is the correlation value of *Z* and *Z′*(2). 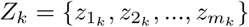 and 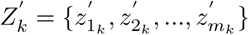 are sets to measure *r*_*IR*_ among items belonging to *X*. The number of items in *Z* and *Z′* is the same. We generated *Z* and *Z′*so that the correlation is close to one. 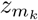 and 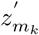 are the each item value of *X*_*k*_. *K* is the number of participants, and *k* is *k*th participant of *K*.

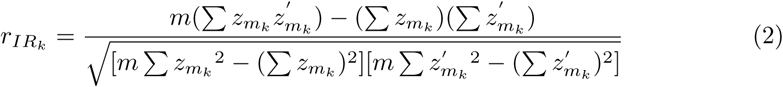

Each index has its cutoff value and calculates the cutoff values for each prepared CRF in a study. In detail, we calculated the mean (*µ*) value and the standard deviation (*σ*) to remove outliers. We removed an outlier if it was greater than *µ* + 3*σ* or smaller than *µ* − 3*σ*. After removing the outliers, we calculated *µ*′ and *σ*′. Then, we calculated the cumulative distribution function (CDF) for the values of DRW indexes (3). The one reason for CDF is to find a cutoff value that has the lowest area under the probability density function among the input values. The other reason is to represent random variables of real values whose distribution is unknown using the collected values [13]. Accordingly, we used the CDF values to find the *µ*″ and *σ*″ again to get a normal distribution. On the distribution, we gained cutoff-value using the z-score for the p-value (default is 0.05). *h*(*X*′) return 0 if the normalized input value is smaller than the cutoff value, and 1 if the value is larger than the cutoff value (4). We calculated the DRW indexes with the binary decision function.

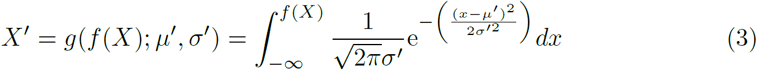

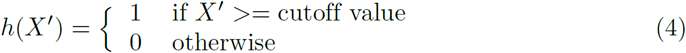

We calculated DRW of a participant when we computed all the cutoff values of DRW indexes for each CRFs (5,6). *N* is the number of CRFs assigned to a participant. 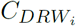 is the number of DRW calculation item counts for each CRF type. Figure 3 displays an example calculation procedure for DRW.

**Figure 3.**
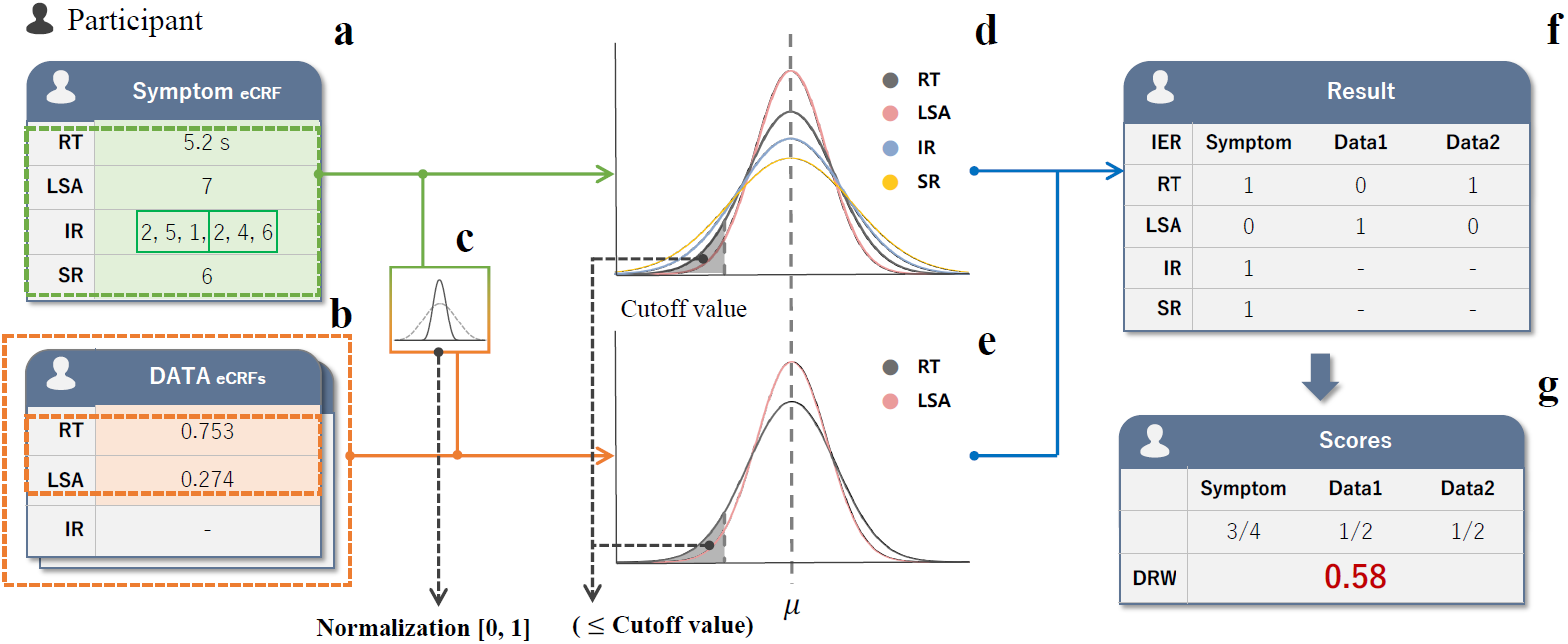
Example of calculating the DRW of one participant in the study. (a) The platform calculates RT, LSA, IR, and SR based on the data entered to calculate the DRW of the symptom of eCRF. Each participant has one symptom eCRF. (b) The platform stores the collected data in data eCRF. Data eCRF, unlike symptom eCRF, does not display a specific pattern, so it calculates RT and LSA excluding IR and SR. (c) The platform calculates the cumulative distribution function for the normalization of input values. (d) The platform calculates cutoff values for RT, LSA, SR, and IR, using values collected from all symptom type eCRFs entered by all participants in a study. (e) The platform groups all participants in the study into an eCRF consisting of the same items of the data type entered. (f) The platform then uses the values collected from each group’s eCRF to calculate the RT, LSA, SR, and IR. (g) The platform calculates DRW index values of the participant using the cutoff values obtained from each eCRF type. The platform uses the obtained cutoff values to calculate the DRW.

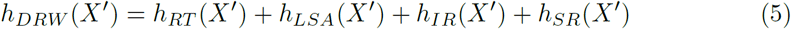

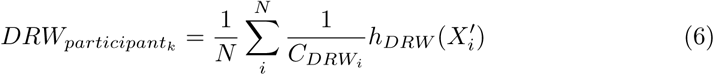

### 4.3 Sustainable Data Sharing

A reward can drive active participation, which is an intuitive fact. Recent scientific findings support the fact that a reward has an impact on data acquisition [29, 49]. We provided financial future rewards for sustainable data collection as exchangeable cryptocurrency [50, 76]. To give the reward, we developed a study result transfer function that delivers statistical information on the completed study to an external cryptocurrency system (Figure 4). The study participants periodically receive cryptocurrency as a reward for their data. The total amount of the reward depends on the external cryptocurrency system. However, the reward is an expected compensation after the end of the study, not an immediate profit. Therefore, we tried to induce sustainable participation with an expectation that active participation will earn high rewards [97]. The expectation established a future reward distribution method to draw continuous management motivation from the creator and the active involvement of participants.

**Figure 4.**
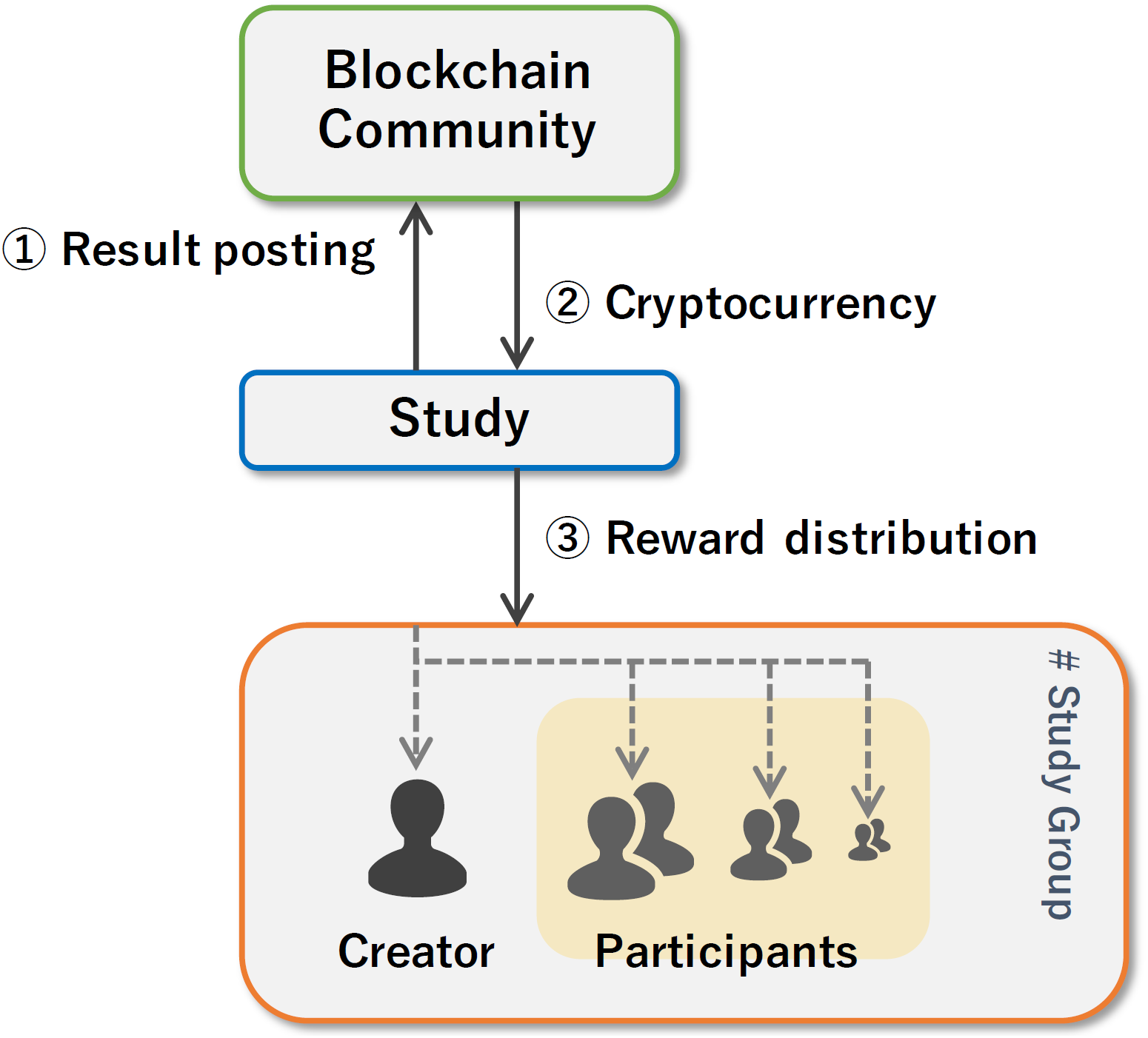
Participation and future rewards of the study. On participating in the study, the result is posted on the blockchain community (➀), and the community provides cryptocurrency to the study (➁). The platform distributes the rewards differently depending on the participant’s level of involvement after the end of the study (➂).

We collect rewards per each completed study. *R*_*total*_ is *R*_*creator*_ + *R*_*participants*_ + *R*_*system*_. *R* are rewards. *R*_*creator*_ calculates the reward amount based on the compensation proportion of participants and *µ*_*DRW*_ value of the study (7). If the *µ*_*DRW*_ is low, this acts as a kind of penalty. We introduced this to encourage the study creators to create a better plan and stimulate a study.

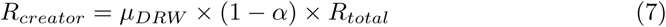

Here, *R*_*participant*_ is that *R*_*total*_ substrates *R*_*creator*_ (8). *K* is the total number of participants. *DRW*_*k*_ is the calculated DRW for *k*th participant. The platform rewards participants for their efforts to input data. A participant receives more rewards if he or she made a more than average effort. Otherwise, the participant gets a lower reward. *R*_*system*_ acquire high compensation when the *µ*_*DRW*_ is low. We added *R*_*system*_ to recover the cost of wasting platform resources when careless participants generated unreliable data. We will use *R*_*system*_ as a platform maintenance cost.

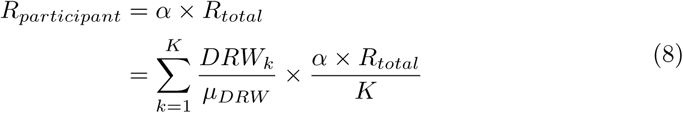

## 5 Pilot Platform Implementation

The proposed platform is designed for a participatory trial. The characteristic of participatory trial in terms of data collection is that people can become researchers and conduct experiments, systematically collecting data based on crowdsourcing and sharing the results with the public [47, 53].

To achieve this goal, we have developed a platform that integrates all the methods presented above, so that each feature is systematically connected to another. The working logic of the integrated platform is implemented as follows (Figure 5). Creators are people who recruit participants to suggest research and evaluate the effectiveness of healthcare apps. Creators build studies in the data preparation phase by using the protocol database and protocol discovery methods. Once the created study is activated, users can join the study. Participants of the study will be curious about the effectiveness of healthcare apps, so that they are more serious about entering data. Research data and basic information about the study are stored in the blockchain to prevent data falsification in vulnerable systems [76]. The study ends after the predefined duration of the research. Even if the number of participants were not enough to develop a sufficient amount of data, the study would be closed at the pre-set endpoint. The system then calculates the effectiveness score from the study data. At the same time, the research results are open to both the public and experts in the data sharing phase, and participants receive rewards based on the data entered.

**Figure 5.**
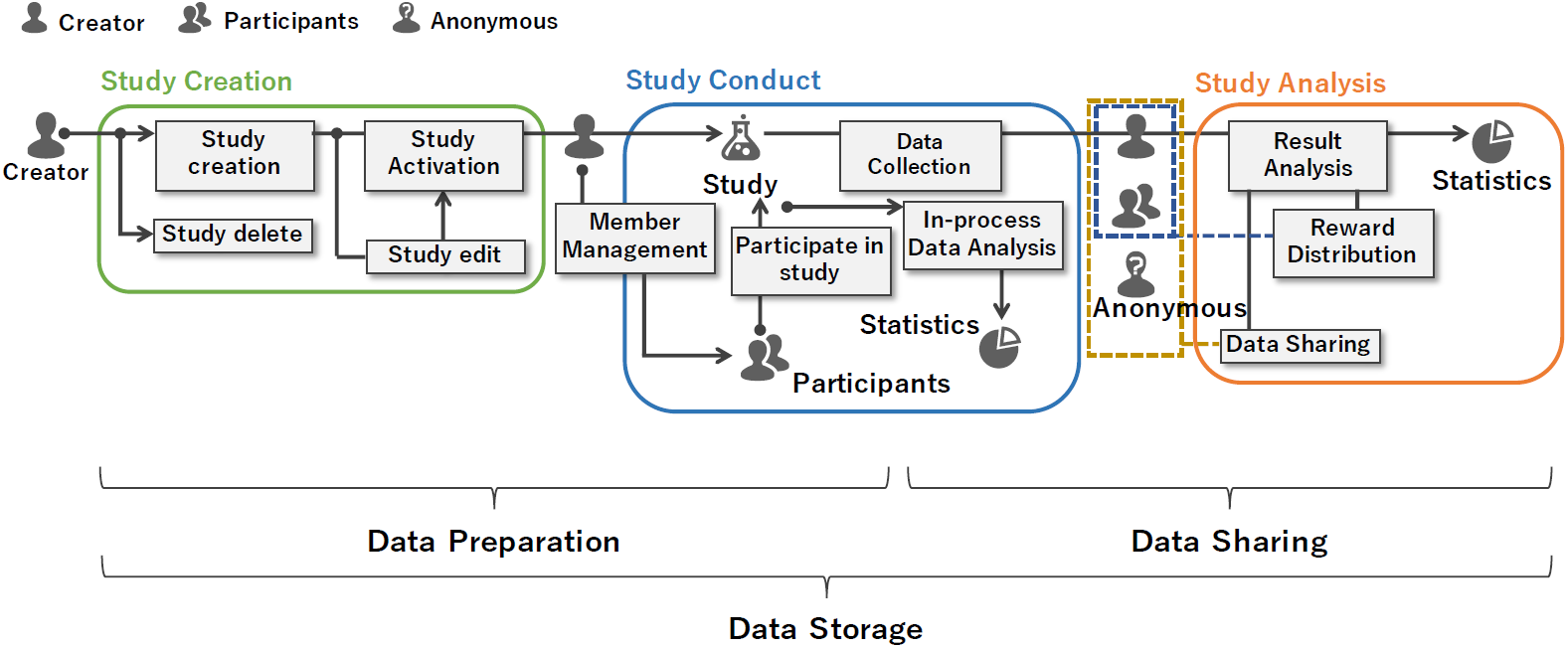
Working logic overview of the platform. The platform integrates and connects methods for reliable data collection at all stages of study creation, study conduct, and study analysis. In the study creation stage, the creator can create, modify, and delete the study. When the creator activates the study, it is open to other participants, and they can participate. In the study conduct stage, the creator can collect the data from the participants and can monitor them. In addition, the creator and participants can monitor statistical information related to the study. In the study analysis phase, all data generated after the study is completed will be disclosed to the study creator, the participants, and the anonymous users. This makes data available to the experts and the public. The platform services allow users to prepare, collect, and distribute reliable data at each stage in the integrated environment.

Technically, we designed the pilot platform according to the interface segregation principle [103], which provides a user with an tool that uses only the services that the user expects. The platform consists of three layers: web interface, API engine, and database. The functions of each layer are based on Node.js, Java programming language, and PostgreSQL database. In addition, essential functions and fundamental database schema related to the EDC system are designed using OpenClinica 3.12.2 [14]. We also inherited the EDC function to receive data in the Operational Data Model (ODM) XML format that is compatible with the Clinical Data Interchange Standard Consortium [10, 33]. Finally, we integrated the blockchain system to prevent data falsification for the crowdsourced data collection platform in participatory trials [76]. See the Supplement document for more information on the platform development.

## 6 Results

The results of the proposed method consist of both quantitative (the test results for each method) and qualitative measurements (a comparison of the platform functions with other platforms). We conducted quantitative measurements with the participants of the platform in terms of simplicity of protocol creation function in the data preparation stage, DRW validation in the data storage stage, and the influence of expected rewards rather than immediate rewards in the data sharing stage. All participants filled and signed an electronic consent form. In the quantitative measurements, the participants’ data input was for the purpose of method validation. We did not collect any personally identifiable information during the tests. We used qualitative measures to demonstrate the benefits of a participatory trial, as compared with clinical and pragmatic trials.

### 6.1 Comparison to the simplicity of data preparation

We developed a protocol retrieval method based on the clinical trial protocol database and validated the method in our previous studies [74, 75]. The methods are the core function of data preparation. In detail, we validated semantic filters of the database that can provide more appropriate protocols than a keyword search [75]. We used clinical trial protocols and corresponding disease conditions as extracted golden standard set from clinicaltrials.gov [107]. Our F-1 score was 0.515, and the score was higher than the keyword search score 0.38. Concept embedding model to find semantic relatedness of clinical trial protocols showed 0.795 Spearman’s rank correlation coefficient with the benchmark set from Mayo Clinic [39, 74]. The result was higher than the previous method Lesk of 0.46 and vector of 0.51. Finally, we conducted a user evaluation test of protocol retrieval system from ten clinical trial experts’ point of view [75]. Our system presented 1.6 difficulties and 6.5 satisfaction using a 7-point Likert scale. Clinicaltrials.gov was 6.2 and 2.3, respectively.

However, previous results were limited by providing tests solely for expert views. Also, we conducted the results for a clinical trial, not a participatory trial. In contrast, the interpretable data preparation method we proposed is a publicly available system to apply the above technique from the perspective of a participatory trial. Therefore, we need further validation of whether the proposed method is sufficiently simple for anyone to create an interpretable protocol to prepare data collection.

For the validation, we designed an experiment that collects Usefulness, Satisfaction, and Ease of Use (USE) score from human participants to reflect their simplicity experience of the different methods [60]. We recruited human participants who represented general users using MTurk [73]. We used 1.1 expected mean and 1.14 expected standard deviation (SD) of the previous USE study [31]. We set a power of 95% and a one-sided level of significance of 5% to calculate the number of participants. The number of participants was 20 by adjusting the sample size for t-distribution [23]. The participants of the experiment compare the data preparation function, which is related to protocol creation between the proposed method and the existing expert method. We prepared the data preparation method as a simulation program (PM), and the complicated method used Openclinica 3.14, an open-source EDC program (OC) [14]. Two programs recorded randomly generated ID of participants to identify their test completion. The participants respond to the USE questionnaire, which consists of 30 items of 7-point Likert scale for the comparison score and 6 short items for their opinion after each use [60]. We reversed the 7-point Likert scale to make participants concentrate more on the questionnaire [11, 41]. We restored the modified scores in the result calculation. In the experiment description, we only provided an elementary data preparation guide and no detailed manual. We also divided participants into two groups to prevent recall bias occurring in the order of the program usage [72]. We reversed the order of use for each group. Finally, we configured the hardware environment identically on the cloud-computing resources of Amazon web services to only identify software differences [17].

After completing the experiment, we excluded participants who did not complete all USE items, had all the same item values, or skipped the test of method systems. Consequently, we could obtain forty-two participants from the combined data set of the two groups. Supplementary data We conducted the descriptive statistics of both methods as four divided dimensions [31]. Accordingly, the 30 USE items broke down as follows: 8 usefulness (UU) items, 11 ease of use (UE) items, 4 ease of learning (UL) items, and 7 satisfaction (US) items. The statistics indicated that the proposed method scores were higher than 5 for all USE dimensions (Figure 6 and Table 2). We also found that scores showed negative skewness for all aspects.

**Figure 6.**
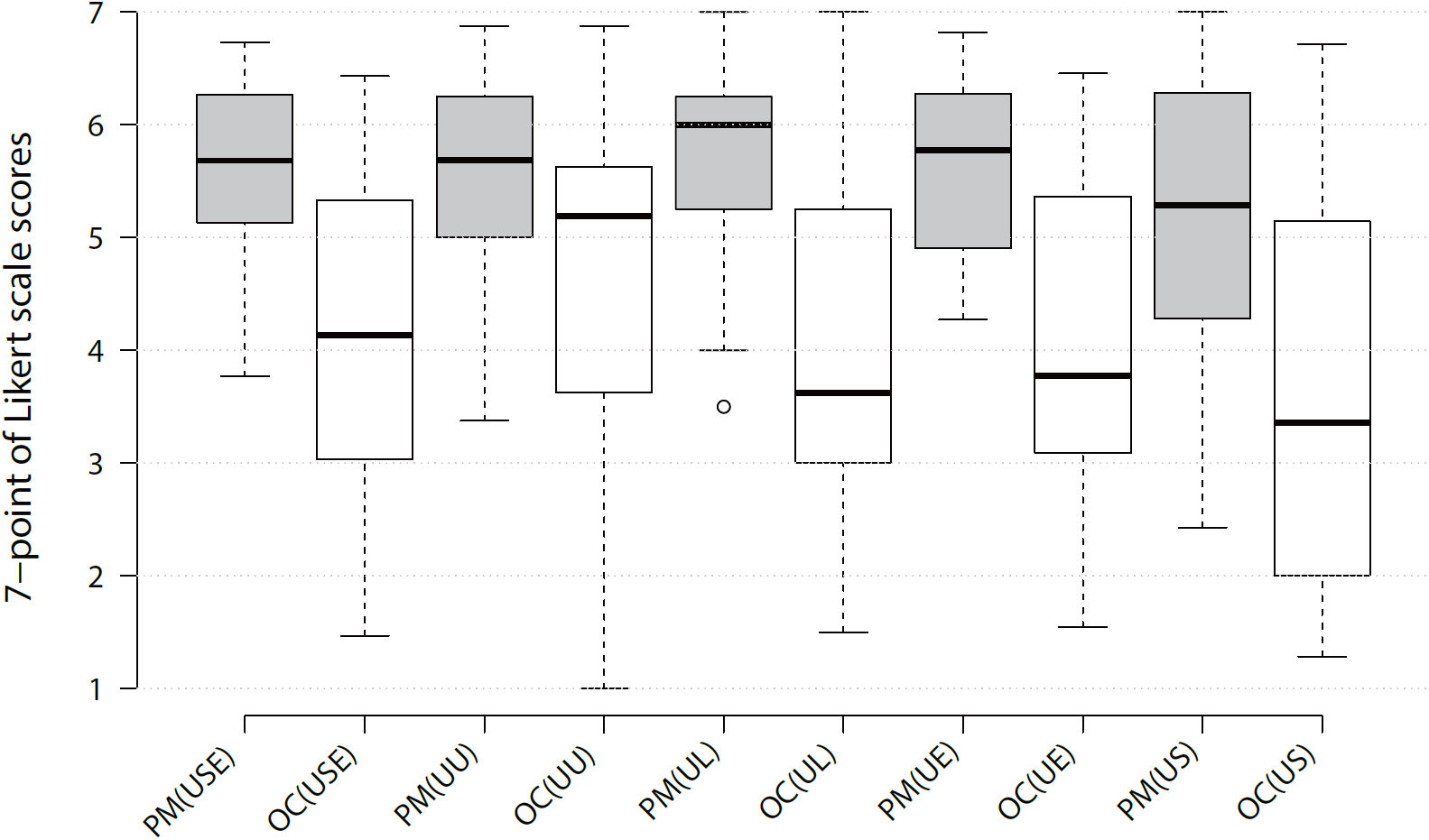
Whisker box plots of the dimensions in USE between two groups. The proposed data preparation method (PM) showed higher records than the existing expert method (OC) in all domains of tests.

**Table 2.** Descriptive statistics of USE in the proposed data preparation method and existing expert method. Statistical description of proposed data preparation method (P) and existing expert method (O).

| **P**<br>O | Mean | Std | Min | Max | Skewness | Kurtosis |
| --- | --- | --- | --- | --- | --- | --- |
| USE | 5.5<br>4.1 | 0.8<br>1.4 | 3.8<br>1.5 | 6.7<br>6.4 | -0.4<br>-0.1 | -0.7<br>-1.0 |
| UU | 5.6<br>4.7 | 0.9<br>1.4 | 3.4<br>1.0 | 6.9<br>6.9 | -0.6<br>-0.7 | -0.3<br>-0.3 |
| UL | 5.8<br>4.0 | 0.8<br>1.5 | 3.5<br>1.5 | 7.0<br>7.0 | -0.9<br>0.2 | 0.1<br>-1.0 |
| UE | 5.6<br>4.0 | 0.7<br>1.4 | 4.3<br>1.5 | 6.8<br>6.5 | -0.3<br>0.0 | -1.2<br>-1.1 |
| US | 5.2<br>3.6 | 1.1<br>1.7 | 2.4<br>1.3 | 7.0<br>6.7 | -0.4<br>0.3 | -0.5<br>-1.3 |

Note. N=42. Min = minimum; Max = maximum

As described in Table 2, the average USE score of PM was 5.548 (SD=0.897), and OC was 4.3 (SD=1.5). We conducted a statistical test to confirm a significant difference between the average scores. To check the USE score’s normality, we did the Shapiro-Wilk test [87]. The test result p-value of our score was 0.082, and the compared score was 0.238, which satisfied normality (p ¿ 0.05). Therefore, we did the paired sample t-tests, and the USE score presented significantly higher value in PM than OC, with t = 7.243, p <=.001. Since the four dimensions in USE did not satisfy the normality, we conducted the Wilcoxon signed-rank test [102]. The outcome of the test showed noticeable differences between the two methods in all dimensions (Table 3).

**Table 3.** Related-samples Wilcoxon signed rank test on the dimensions in USE between proposed data prepration and existing expert method

| Dimension | N | s.t.s. | p |
| --- | --- | --- | --- |
| Usefulness (UU) | 42 | -4.327 | $\leq .001$ |
| Ease of Use (UE) | 42 | -5.196 | $\leq .001$ |
| Ease of Learning (UL) | 42 | -5.125 | $\leq .001$ |
| Satisfaction (US) | 42 | -4.848 | $\leq .001$ |

Note. s.t.s = standardized test statistic

Besides the results, we also found that the participants completed 100% of tasks on PM, while they only generated basic metadata of given tasks on OC. Thus, we concluded that PM is more suitable for understandable data preparation in participatory trials.

### 6.2 Data Reliability Weight Validation

DRW is a weight that is calculated by incorporating a score to measure the effort spent when a user enters data. We examined the correlation between the Human Intelligent Task Approve Rate (HITAR) of MTurk and DRW to assess whether this score is measured well. The HITAR is a score that evaluates how well a worker is performing tasks assigned by the requestor on MTurk. Consequently, we assumed that a higher HITAR would lead to a higher DRW. To verify this, we performed the following tests.

We prepared the first test to confirm that the DRW indexes correlated to HITAR. DRW indexes include response time (RT), long string analysis (LSA), individual reliability (IR), and a self-report (SR) [41, 42]. We intended to organize DRW with empirical DRW indexes for simple calculation in eCRF. Because DRW is a subset of IER, we made a questionnaire to calculate DRW based on detecting IER research [41]. We prepared a questionnaire to contain 60 items of the international personality item pool—neuroticism, extroversion, and openness (IPIP-NEO)—to calculate IR [45].We also added 8 items of the infrequency IER scale to increase the concentration of participants at the beginning of the test and SR item for DRW score [65, 77]. We placed a self-input HITAR item box at the start of the questionnaire as LSA item. Thus, the questionnaire included 70 items. To calculate the total sample size, we used 0.13 expected mean of the paired differences based on the result of our pilot DRW test with a total of 100 participants in MTurk. We set a power of 95%, a one-sided level of significance of 5%, and equal group size for sample size calculation [23]. The calculated sample size was 153 for each group, and the total size was 306. We recruited the participants using MTurk. Workers of MTurk participated in the study as participants. We divided all participants into two groups, with HITAR levels above 95 and below 90. Participants over 95 HITAR needed a record of more than 500 tasks before participating. This was to select a participant with an average of more than one year from the average number of 1,302 tasks of approximately 2.5 years analyzed in the previous study [37]. Participants below 90 HITAR had no other restrictions. This is because the first granted HITAR is 100, and it is a deduction method when the task is not performed well. Participants read the task description posted on MTurk. All settings of the test were identical except for the HITAR levels. The endpoint of the test was set to the recruitment completion time of one of the groups.

A total of 340 participants were involved in the test (Supplementary data 2). Each group had 170 participants, and we considered that the number of participants reflected the calculated sample size. Based on the collected data from the participants, we compared the average value of DRW and DRW indexes per each group. In all DRW indexes and DRW, groups with more than 95 HITAR scored higher than the other group (Figure 7 (a)). In more detail, we did a statistical analysis to understand the difference between the two groups. Table 4 displays the statistical description. The DRW index values in the table 4 were converted to CDF value to calculate the DRW. SD in all areas except RT and LSA showed similar values. The reason was that 95 HITAR group had very high values in RT and LSA. We calculated cutoff values using CDF values through DRW calculation method. We checked that only passes a range of values with approximately 0.5 or more. We verified statistical significance between the mean values of the two groups using the independent samples t-test. We performed Levene’s test to consider unequal variance in the t-test [58]. The p-values indicate that the results are statistically meaningful. Only SR had similar values in the two groups. We presumed that the mean value difference of SR was small, but SR would have a small effect on the calculation of DRW. Furthermore, we calculated effect size, an indicator of how much difference there is between the two groups [64]. Since we are comparing the same sample size between the two groups, we calculated the effect size using Cohen’s d, and the result was close to a large effect (.8) according to the interpretations of Cohen’s d [18].

**Table 4.** Statistical description and independent samples t-test of IR, LSA, RT, SR, DRW between groups of HITAR values greater than 95 and less than 90.

| Index | HITAR condition | Number of participant | Mean | Standard deviation | Cutoff value | t | d.f. (degree of freedom) | p | Cohen's d |
| --- | --- | --- | --- | --- | --- | --- | --- | --- | --- |
| DRW | >= 95 | 170 | 0.582 | 0.232 | N/A<sup>a</sup> | 6.009<sup>b</sup> | 338 | <.001 | 0.652 |
|  | <= 90 |  | 0.425 | 0.251 |  |  |  |  |  |
| IR<sup>c</sup> | >= 95 |  | 0.460 | 0.272 | 0.418 | 2.310 | 338 | 0.021 | 0.251 |
|  | <= 90 |  | 0.391 | 0.276 |  |  |  |  |  |
| LSA<sup>c</sup> | >= 95 |  | 0.553 | 0.193 | 0.473 | 6.690<sup>b</sup> | 294.743 | <=.001 | 0.726 |
|  | <= 90 |  | 0.375 | 0.289 |  |  |  |  |  |
| SR<sup>c</sup> | >= 95 |  | 0.555 | 0.254 | 0.577 | 0.723 | 338 | 0.470 | 0.078 |
|  | <= 90 |  | 0.535 | 0.255 |  |  |  |  |  |
| RT<sup>c</sup> | >= 95 |  | 0.518 | 0.263 | 0.452 | 6.987<sup>b</sup> | 263.457 | <=.001 | 0.758 |
|  | <= 90 |  | 0.357 | 0.146 |  |  |  |  |  |

*Note.* <sup>a</sup>Not applicable <sup>b</sup>The t-test result was reported on the unequal variances based on the Levene's test. <sup>c</sup>The value is normalized by CDF.

**Figure 7.**
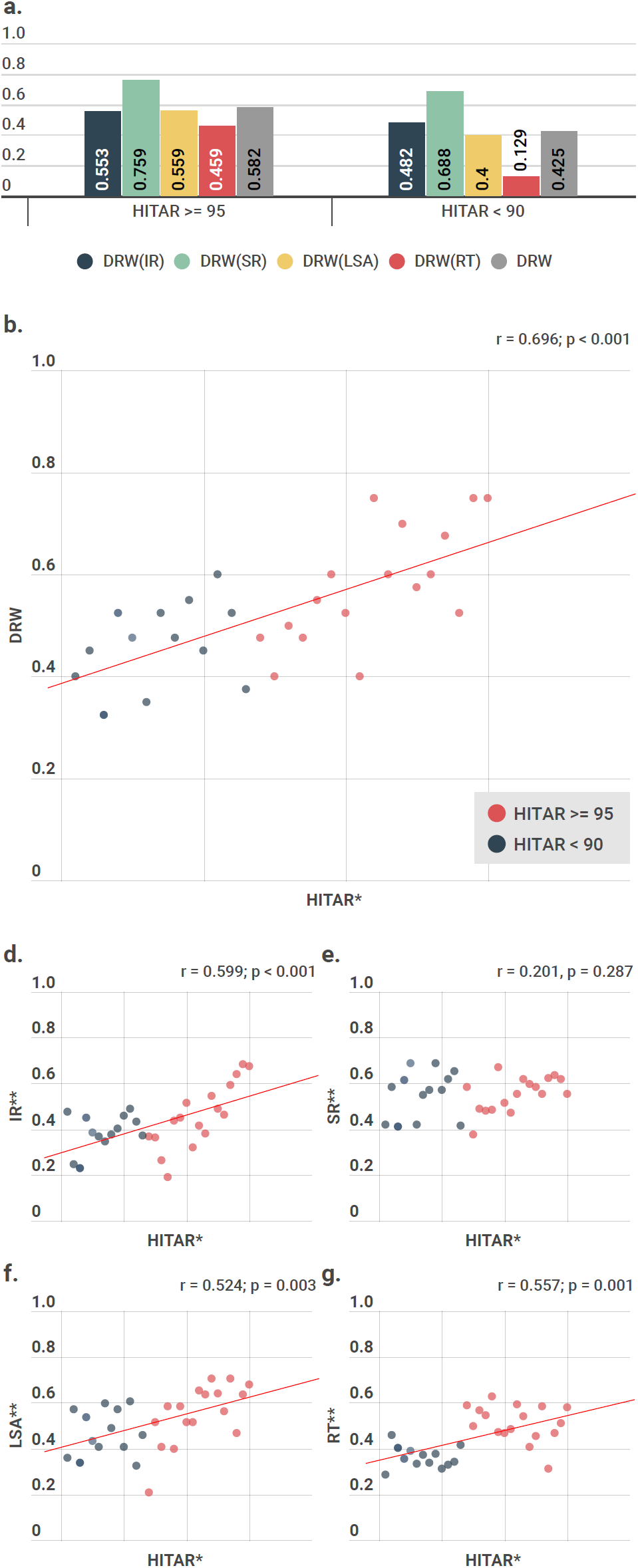
Correlation between DRW and HITAR. (a) Average DRW indexes of worker groups classified by HITAR; (b) correlation between HITAR and DRW; (d-g) correlation between HITAR and DRW indexes; * is scaled values. ** is normalized values.

Next, we compared the correlation between DRW values and HITAR. The obtained HITAR values presented high skewness (−2.138) because the HITAR average we obtained had a relatively high value of 83.91. To improve the performance of the data analysis and interpretation, we established a binning strategy. We thus divided the data group by the same frequency, and we smoothed the grouped data values to the average value. Specifically, the actual data processing is as follows. We sorted the number of 340 data of participants based on descending order of HITAR values. We grouped 10 as frequency and averaged the HITAR, DRW, IR, SR, LSA and RT values. Based on the groups, we ranked HITAR and took the reverse order. 34 Participants did not enter HITAR values correctly, and 6 data of participants remained as ungrouped. Accordingly, we removed the 40 data, and we placed 300 data into 30 groups. To check the effectiveness of the binning, we used an information value (IV) that expressed the overall predictive power [86]. The measured IV value (0.465) was greater than 0.3, and we determined that the binning was good. Based on the binning, collected data showed significant results that indicated a noticeable correlation between DRW and HITAR (Figure 7 (b)-(g)). DRW and HITAR also presented an interesting correlation score (r=0.696, p ¡ 0.001). In summary, we validated that HITAR correlates with DRW and DRW can be used as a reliability measure.

### 6.3 Future Reward Distribution Effect Observation

Recent studies have confirmed that systems with immediate virtual rewards have a positive effect on continuous data collection [68, 78]. However, we evaluated the validity of a reward distribution that gives rewards to participants in the future, rather than immediately. We showed the validity as correlation between rewards and DRW. Accordingly, we evaluated the following scenario, in which the future reward system of our platform increases the participation rate.

We created a simulation environment in MTurk to reduce the observation time in the real world. We agreed that a real-world study of the reward distribution effects would require significant time to collect the data and cases. For example, PatientsLikeMe required approximately 5 years to collect enough cases to show the benefits to communities (which were related to the extent of site use) [101]. Thus, we conducted two tests in the simulation environment. We designed one test that informed about reward distributions in the future (the same as our platform). The other test did not contain this information. The remaining settings were unchanged. For the sample size calculation, we conducted a pilot test with 100 participants in MTurk. We obtained 0.11 expected mean of the paired differences and 0.25 SD on the result of the pilot test. Based on the result, we calculated the sample size as 168 per each group to have a power of 95% and a one-sided level of significance of 5% [23]. The total sample size was 336. Then, we used a questionnaire as data reliability validation and weights for consistency comparison, except for the HITAR. However, we needed to consider the casual observation characteristics of MTurk workers since this experiment had no constraint on HITAR [5]. Therefore, we added five questions that induced to enter as many words as possible in the LSA index of DRW [42].

We recruited 336 workers from MTurk to participate in the test. We allocated 168 workers to the test that contained future reward information (RI) and the other 168 to the test without future reward information (NRI). We designed the test to be completed within two hours, and we recorded the test start and end times for each worker as RT. Consequently, we successfully collected data from 336 workers (Supplementary data 3). First of all, we analyzed the effect of RI on the self-reporting rate by a two-way ANOVA test. Table 6 indicates that the reward condition has interaction with consecutive LSA items and is statistically significant at an alpha level of 0.05 (p = 0.008). In other words, we found that continuously containing long characters in consecutive LSA items on the RI condition, and we inferred that the RI condition improves the rate of self-reporting than NRI condition. Next, we found that the average DRW of RI was 0.605, and the DRW of NRI was 0.472 (Figure 8). Table 5 shows the significant difference between DRW of RI and NRI (p=0.03). Figure 8 presents that DRW(SR), DRW(LSA), and DRW(RT) of RI also have higher average values than the DRW indexes of NRI, and Table 5 provides detailed statistical information. DRW index values in Table 5 are CDF converted values to calculate the DRW. The DRW(IR) showed a low average result in RI, but we interpreted that the effect size (0.076) of the index is too low to effect the DRW result in Table 5. We presumed that this is caused by not controlling workers with HITAR. Interesting cases are found in both RT and LSA. Both indexes showed high average DRW values and a significance difference in Table 5. In particular, we configured the DRW method to measure RT and LSA only in the data type CRF, and we could confirm that the configuration was appropriate according to the effect of rewards on continuous data collection. Thus, we concluded that the reward distribution not only increased DRW but also helped to improve the sustainability of the data collection platform.

**Table 5.**
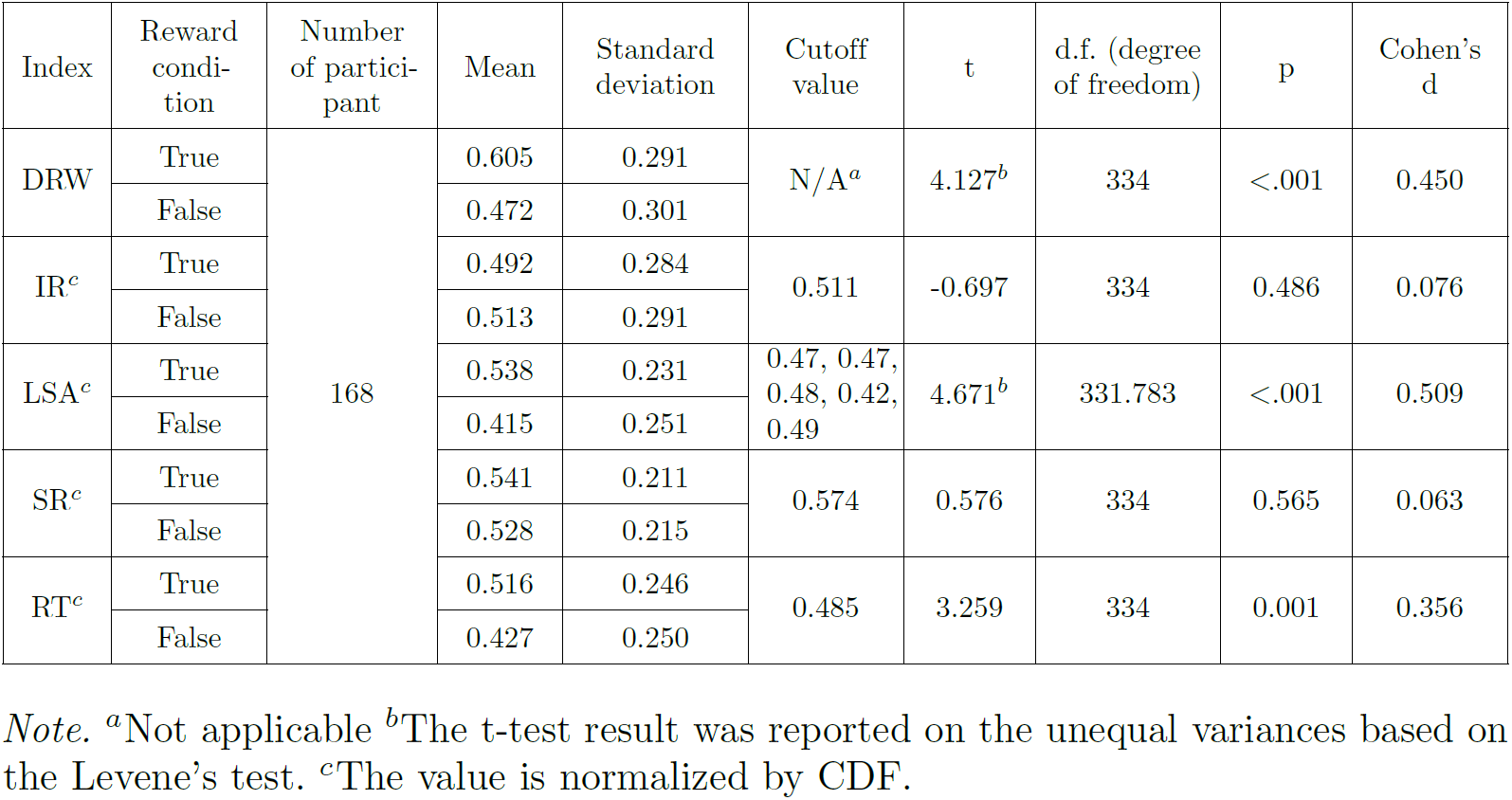
Statistical description and independent samples t-test of IR, LSA, RT, SR, DRW between groups of future reward and no future reward information.

| Index | Reward condition | Number of participant | Mean | Standard deviation | Cutoff value | t | d.f. (degree of freedom) | p | Cohen's d |
| --- | --- | --- | --- | --- | --- | --- | --- | --- | --- |
| DRW | True | 168 | 0.605 | 0.291 | N/A<sup>a</sup> | 4.127<sup>b</sup> | 334 | <.001 | 0.450 |
|  | False |  | 0.472 | 0.301 |  |  |  |  |  |
| IR<sup>c</sup> | True |  | 0.492 | 0.284 | 0.511 | -0.697 | 334 | 0.486 | 0.076 |
|  | False |  | 0.513 | 0.291 |  |  |  |  |  |
| LSA<sup>c</sup> | True |  | 0.538 | 0.231 | 0.47, 0.47, 0.48, 0.42, 0.49 | 4.671<sup>b</sup> | 331.783 | <.001 | 0.509 |
|  | False |  | 0.415 | 0.251 |  |  |  |  |  |
| SR<sup>c</sup> | True |  | 0.541 | 0.211 | 0.574 | 0.576 | 334 | 0.565 | 0.063 |
|  | False |  | 0.528 | 0.215 |  |  |  |  |  |
| RT<sup>c</sup> | True |  | 0.516 | 0.246 | 0.485 | 3.259 | 334 | 0.001 | 0.356 |
|  | False |  | 0.427 | 0.250 |  |  |  |  |  |

*Note.* <sup>a</sup>Not applicable <sup>b</sup>The t-test result was reported on the unequal variances based on the Levene's test. <sup>c</sup>The value is normalized by CDF.

**Table 6.**
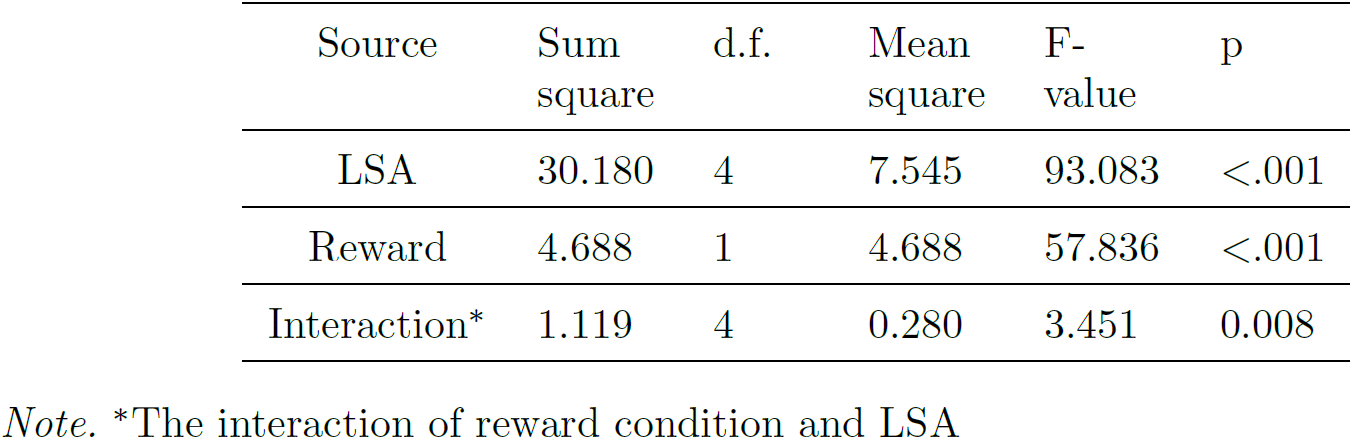
The two-way ANOVA test considering the interaction of reward condition and LSA

| Source | Sum square | d.f. | Mean square | F-value | p |
| --- | --- | --- | --- | --- | --- |
| LSA | 30.180 | 4 | 7.545 | 93.083 | <.001 |
| Reward | 4.688 | 1 | 4.688 | 57.836 | <.001 |
| Interaction* | 1.119 | 4 | 0.280 | 3.451 | 0.008 |

*Note.* \*The interaction of reward condition and LSA

**Figure 8.**
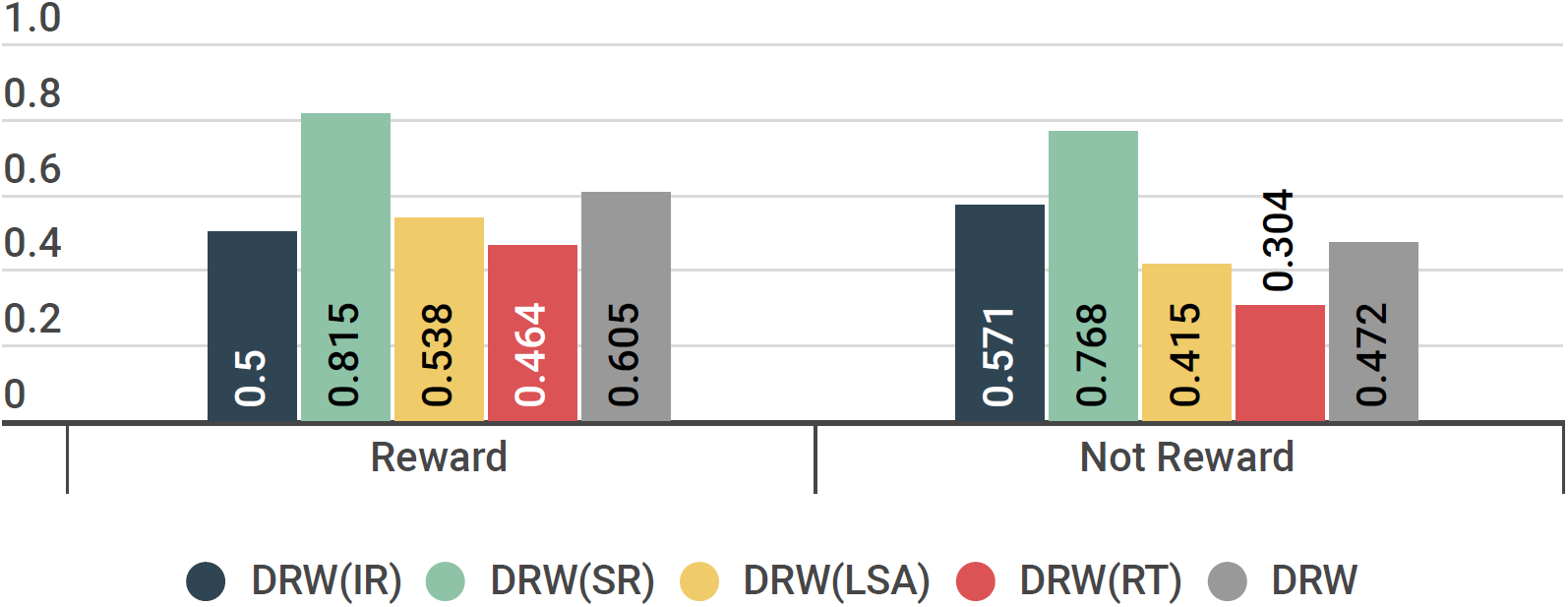
Reward distribution effects. Each average score of DRW index for all workers per test.

### 6.4 Overall Platform Function comparison

We developed “Collaborative research for us” (CORUS, https://corus.kaist.edu) as a pilot data collection platform for participatory trials. CORUS is designed based on the EDC for existing clinical trials, but it makes it easier to design a study and provides services that allow users to participate voluntarily and continuously in the data collection for the study. In this section, we compare and analyze the features of CORUS in detail. For specific comparisons, we selected the programs mentioned or used in journals related to clinical research from 2012 to 2019, and we introduced them in the introduction section. We summarize the comparison results in Table 7 and provide the details in the following subsections.

**Table 7.**
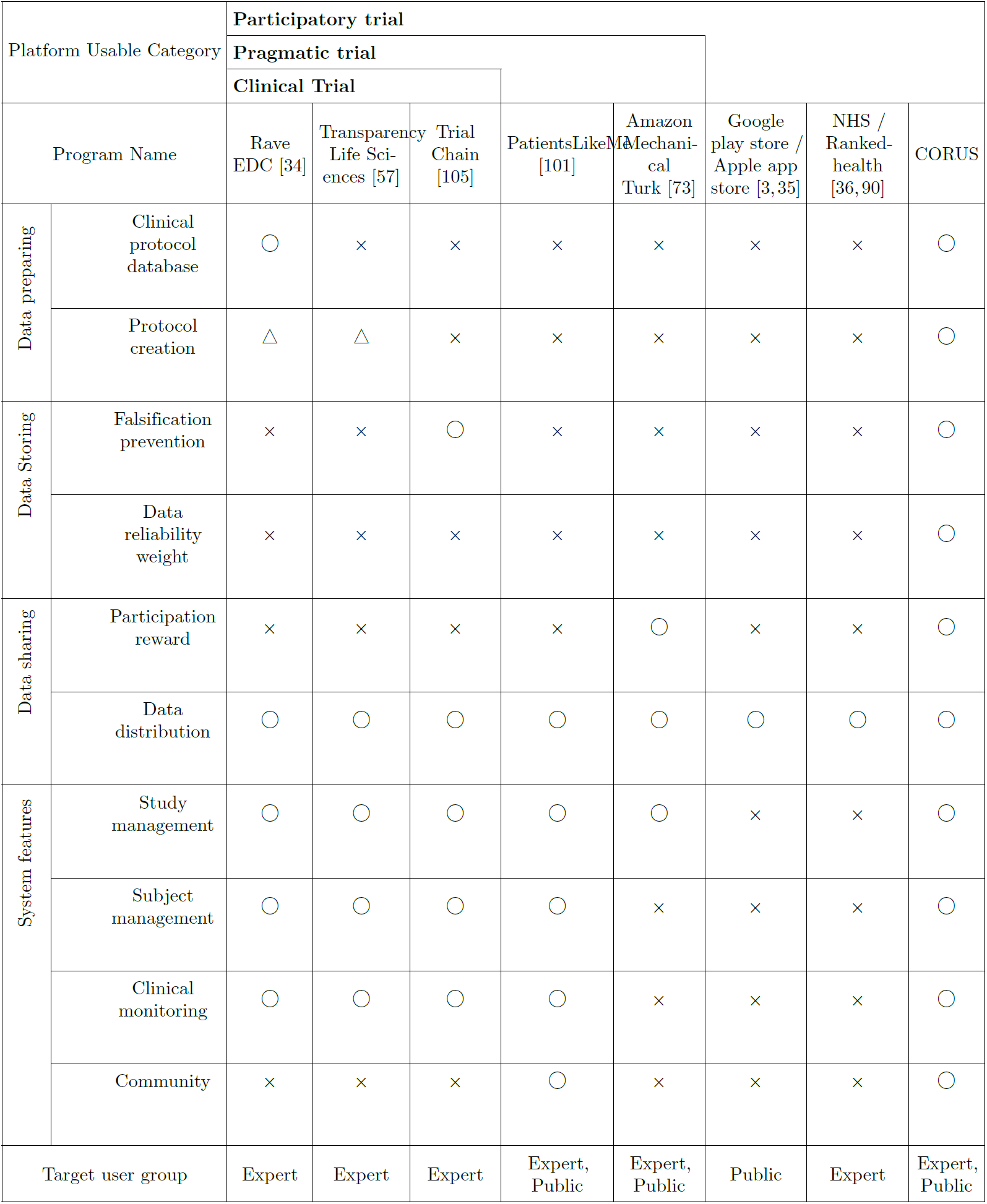
Platform function and feature comparison in a manner of reliable data collection for the participatory trial

#### 6.4.1 Comparison with the clinical trial systems

CORUS has a protocol creation feature as interpretable data preparation that makes it easy for the public to use without the help of experts. In the platform, the creator can utilize the clinical trial protocol database feature according to the semantically related symptom of the effectiveness of healthcare apps. Moreover, the creator can get feedback from the study participants within the community feature, which allows the creator to enhance the protocol in the future.

#### 6.4.2 Comparison with the pragmatic trial systems

The protocol creation feature of CORUS provides understandable and straightforward data preparation for public users. The system features of CORUS helps the users lead a study without expert knowledge. Moreover, CORUS has the DRW feature that automatically calculates the reliability scores collected data for reliable data collection. The scores are an objective value that does not comprise the subjective intention of the study creator. We also developed the scores to evaluate the effectiveness score of apps in further study of data storage phase.

#### 6.4.3 Comparison with participatory trials

We developed CORUS as a participatory trial platform. Data preparation features of CORUS prepare data collection that can perform immediate data analysis. Postprocessing for unformatted data is not necessary on the features. The features minimize the flawed study design bias [72]. Besides DRW, data falsification prevention feature prevents transfer bias by not exposing the collected data to the participants during a study. CORUS, a participant-driven platform, also supports additional analysis for experts. Standardized data distribution is an essential feature for validating the effectiveness of an app. Cryptocurrency is a distinctive participation reward feature of CORUS. We connected CORUS to an external blockchain to encourage participants to share their data continuously. Participants can earn a future reward based on their data. CORUS calculate the participant portion of total cryptocurrency of a study based on DRW of the participant. Thus, the participation reward can induce continuous engagement and reliability.

A common limitation of all mentioned platform, except CORUS, is the data falsification problem. The falsification possibility of the collected data, whether intentionally or unintentionally during a study, exists in all platforms [32]. Trialchain used blockchain to solve the problem [105]. We classified Trialchain as EDC based on the explanation of the platform. Thus, we considered that Trialchain is difficult to use in a participatory trial. CORUS also uses Blockchain technology. We developed a data falsification prevention feature based on the data immutability characteristics of blockchain technology [76]. The feature solves the problem of data falsification.

## 7 Case Study: operational test of the pilot platfom

We conducted an operational test to observe the data collecting capabilities of the platform in actual participatory trial conditions. The objective of the test was to confirm whether an app that enables blue light filters on smartphones would help to improve sleep disorders. Although there have been previous studies that showed the relation between blocking blue light and improvement of certain sleep disorders, to the best of our knowledge, it has not been clinically confirmed whether the blue light filters on smartphones or other mobile devices have the same effect [27]. The test was designed to recruit 100 participants. Based on the collected data, we could not find the effectiveness evidence for the app.

We designed the participatory trial project and posted it on the test platform. In the trial, participants were asked to apply the blue light filters (which blocked blue-wavelength lights on their smartphones) and report changes in the quality of their sleep. When users selected a specific project on the platform, introductions to the projects by the creators were shown. This made it convenient for users to participate in their preferred projects. Participants enrolled in the project could immediately participate in the trial and conveniently report data. The platform also encouraged participants to report their data daily by using features such as leaderboards and alarms.

In addition to technical validation, we also validated whether the system could effectively recruit participants to the trial. The duration of participant recruitment was originally scheduled for a month, but the actual rate of recruitment was faster than that. In total, 100 participants were recruited in 21 days. We compared the rate of recruitment in this experiment with those known from previous studies. To denote the rate of recruitment among the trials, the recruitment rate was defined as the number of participants recruited per month (or participants per center per month, in the case of multicenter trials). In 151 traditional clinical trials supported by the United Kingdom’s Health Technology Assessment program, the recruitment rate was 0.92 [98]. According to 8 web-based or mobile app-based studies collected from literature databases, an average of 468 participants was recruited over 5 months (recruitment rate = 93.6) [54]. For the platform, the recruitment rate was 142.8. This shows that participant recruitment using the platform was significantly more effective than participant recruitment in traditional clinical trials; it was also competitive with other web-based or mobile app-based studies. At this stage, which was the beginning of the study period, the response rate also tended to increase over time. The response rate increased to 120% on recruitment days 17–18, when many new participants were enrolled, and the response rate remained high for a significant period of time (Figure 9).

**Figure 9.**
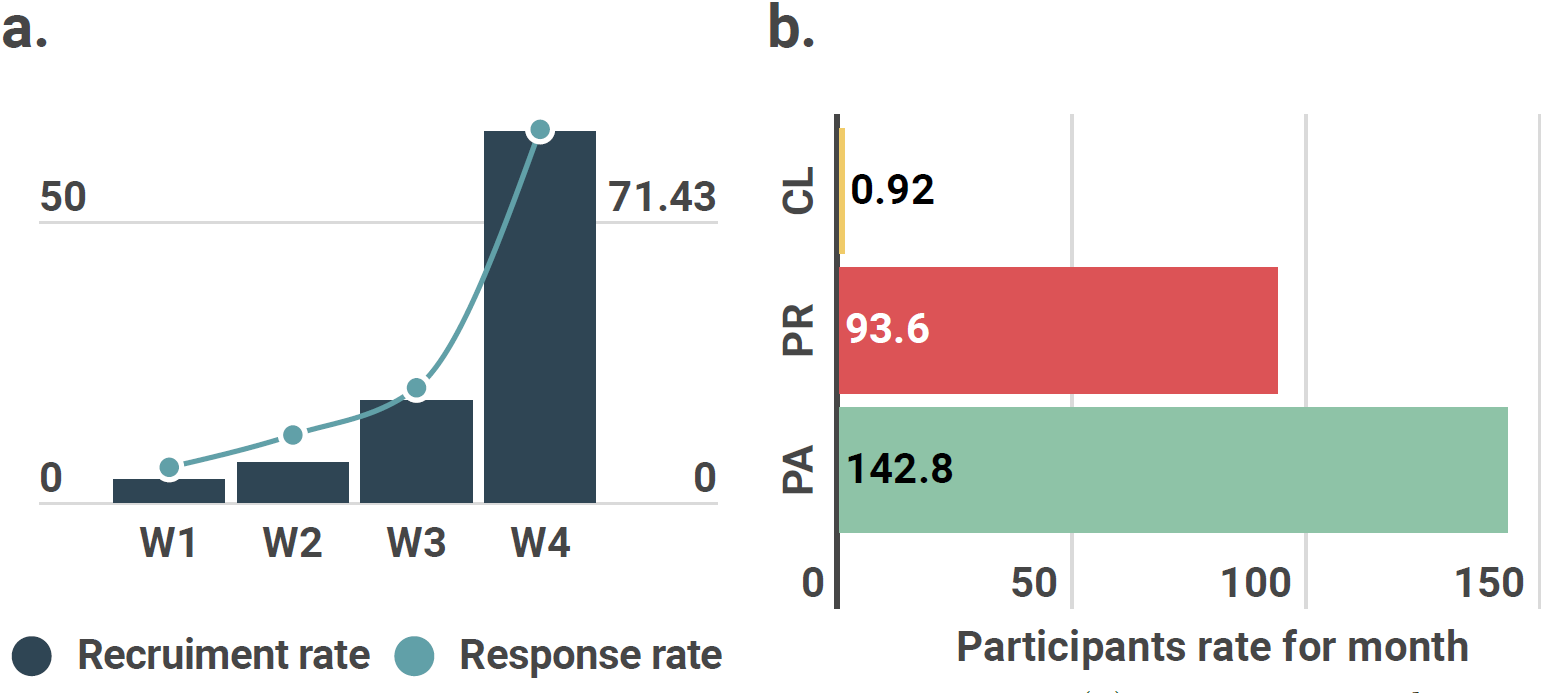
Recruitment rate and comparison analysis. (a) recruitment and response rate for a month from a platform test study; (b) participant rates among clinical trial (CL), pragmatic trial (PR), and participant trial (PA) based on the literature evidence

## 8 Discussion

Our method and platform have technical value in combining various independent and unrelated methods. We integrated a clinical trial protocol and database, semantic correlation calculations, blockchain, survey data evaluation, intrinsic and extrinsic motivation of human behavior, cloud computing technology, and software engineering methods into a participatory trial platform to assess the effectiveness of healthcare apps. Although there are many types of participatory trials, such as crowdsourcing and crowd-science, these approaches still have critical limitations in unstructured data collection and data sharing. We suggested a new concept, which is a participatory trial, to combine beneficial characteristics of crowd-science and crowdsourcing. Then, we proposed a solution to solve problems on interpretable data preparation, systematic data storage, and sustainable data sharing stages. Our method and platform have great value in presenting a primary technique that can validate the effectiveness of various healthcare apps.

Minimizing bias occurred intentionally or unintentionally at all stages of data collection is another important factor in ensuring data reliability. We identified and considered each source of bias in the data collection stages according to relevant research [72, 89]. At the stage of data preparation, we included flawed study design, channelling bias and selection bias. We tried to prevent flawed study design by using the existing clinical trial protocols. We minimized channelling bias, which leads to including only specific participants, by placing unconditional eligible criteria that anyone can participate in a study. The editable eligible criteria of a creator such as gender and age is still a matter to consider for the channelling bias. However, the selection bias, which can distort the data tendency, is not considered. This is a difficult problem to solve in the field of web-based data collection [7]. As a solution, we propose the social relationship distance estimation method to keep participants beyond a certain distance. A social network estimation method of participants to calculate a closeness rank among participants is an example method for cutting off a certain distance or less [82]. At the stage of data storage and data sharing, we included interviewer bias and transfer bias. The interviewer bias refers to potential influence on the participants deriving from the data collector. Our data storage method does only involve mechanical data collection functions, so we assumed interviewer bias not to occur. However, we do not have any preventive measure against interviewer bias caused by the study description or the CRF items that a creator manually generates. The solution is to clearly define the digital biomarkers that a creator seeks to evaluate the effectiveness of a digital healthcare app [20]. Further research and consensus from experts are needed to enable it for standardized participatory trial protocols with digital biomarkers. The transfer bias in our method is caused by data exposure at the data storage stage and by possible influence by the stored data on other participants. We prevented this source of bias by preventing the stored data from being exposed during a study. Finally, we did not consider the bias generated at the stage of data analysis because it is out of the scope of this article, which is only dealing with data collection methods.

There are additional considerations that may improve our platform. First, the current platform cannot provide a quantitative comparison of the search results of the clinical trial protocol. The platform takes a symptom from the creator and provides all relevant clinical trials. However, many clinical trials compare as a search result, additional analysis from the creator’s side is required. In clinical trials, various types of data—not only symptoms but also genes and chemical compounds—come up. This information can be used to retrieve clinical trial data required by the user. If an advanced platform is developed in the future that can quantitatively assess the relevance of all data involved in the clinical trial, it will reduce the additional analysis burden for the creator. Second, advanced DRW methods should be developed. For example, the basic DRW can be used to assess the reliability of the input data based only on the length of the input text, as in long-string analysis (LSA). In other words, it cannot consider semantic errors in the input text. Therefore, an advanced method that considers semantic errors in the text is required. In addition, we have a cutoff value calculation problem when the SD is 0 from the collected data of a DRW index. We prevented the problem with a predefined minimum cutoff value on the platform, but we need to improve this approach so that we can actively find cutoff values suitable to the collected data. Third, we will devise an optimal strategy to reduce the cost of running the platform. Fourth, an additional study is required to verify the platform. We plan to conduct a small-scale pilot study to check whether the study results are the same as the verified effects.

All these limitations should be considered for further experiments or improved computational methods. Moreover, we suggest further studies since the data collection methods are the basics of the digital healthcare field, which is under tremendous progress. The suggested studies are the following: (1) Developing a standard for a participatory trial protocol method considering digital markers and confounding factors, (2) Advancing protocol creation methods for the determination of specific parameters, such as the number of uses, effective report counts, study duration or the number of participants, (3) Using natural language question models to generate DRW index questions rather than human-managed templates, (4) Optimizing data reliability weight calculation with the importance value of each index, (5) Implementing an effectiveness score calculation method based on the collected data with DRW scores. With these further improvements and further studies, our platform could be used as a valuable tool to assess the effectiveness of various apps in a cost-effective manner.

## 9 conclusion

At the first stage of the verification workflow, we proposed the reliable data collection method and platform to assess the effectiveness of digital healthcare apps. We presented a participatory trial concept for a new type of data collection trial that uses voluntary participation of end-users of digital healthcare apps. Then, we described the essential methods of the reliable data collection platform based on the participatory trial concept. Interpretable data preparation methods consisted of a protocol database and a retrieval system to create a protocol without expert knowledge. We validated the simplicity of the methods from compared USE scores between the proposed system and the existing system. We developed a DRW calculation method for systematic data storage. The DRW score showed to be a reliable measure through correlation with HITAR. To achieve sustainable data collection and sharing, we developed a reward distribution method. We indirectly observed the effect of reward using DRW. The effect of reward presented increasing DRW, i.e., it increased the participants’ effort toward sustainable data collection. Finally, we implemented a pilot platform, CORUS. CORUS integrated all the methods and essential features for a participatory trial platform. We compared its features with existing platforms in the field of clinical trial, pragmatic trial and participatory trial. We assert that CORUS has all the necessary features to collect reliable data from the users of digital healthcare apps on the path of validation of their effectiveness.

## Code Availability

We provide all the source code at https://github.com/junseokpark/corus upon the publication. The platform is open-source to promote development in the public domain. The repository describes software requirements and distributes a README.MD file.

## Acknowledgment

Thanks to Jooyeon Lee and Inhwan Hwang of Amazon Web Services Korea LLC (AWS Korea) Worldwide Public Sector for their support with cloud computing platform services and demonstrations.

