## Supplement document for "Reliable data collection in participatory trials to assess digital healthcare apps"

#### Database

http://d2jjojlo5w7xa7.cloudfront.net/database_capture.pdf

**Supplementary Figure 1. CORUS's Main Table Entity Relation Diagram** The platform database consists of tables related to Study, Event, CRF, Item, and Subject. One study is created based on the table structure. Event, CRF, and Item are created accordingly, and related data is added according to the subject's participation. As of November 15, 2019, the CORUS DB consists of 170 tables and 41 views, but this structure might change due to ongoing development requirements. More details on this can be found at the previously mentioned GitHub address and you can download and use the schema structure file.

**Supplementary Table 1.** eCRF types and their description

| CRF Type ID | Name | Description |
|---|---|---|
| 1 | SYMPTOM | Symptom investigation of a participant |
| 2 | ELIGIBILITY | Eligibility checks to participate in a study |
| 3 | CONSENT | Voluntary consent to participate in a study |
| 4 | DATA | Data collected to find effectiveness of healthcare apps |

**Supplementary Table 2.** Item data types for eCRF

| Item_data_type_id | Code | Name |
|---|---|---|
| 1 | BL | Boolean |
| 2 | BN | BooleanNonNull |
| 3 | ED | Encapsulated data |
| 4 | TEL | Telecommunication address |
| 5 | ST | Character string |
| 6 | INT | Integer |
| 7 | REAL | Floating |
| 8 | SET | Group values |
| 9 | DATE | Date |
| 10 | PDATE | Partial date |
| 11 | FILE | File |

#### Service architecture

We categorized this pilot program as a platform because it could be the basis for external applications beyond its original purpose. We designed a working logic to integrate all methods mentioned above: data preparation, data storage, and data sharing (Supplementary Table 3). We chose Amazon Cloud as environment for this platform. Moreover, we designed our server architecture using Amazon services to create a highly available, stable, and secure system environment (Supplementary Figure 2).

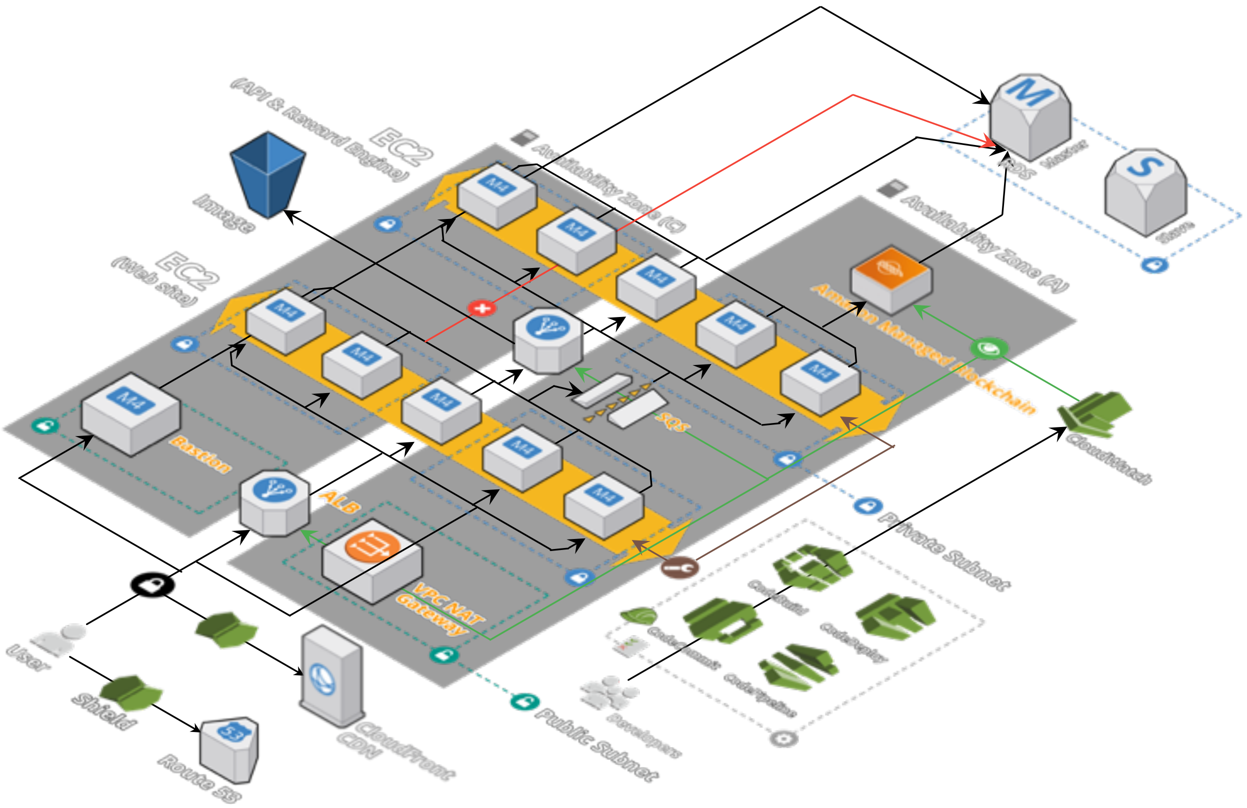


**Supplementary Figure 2. Overall server architecture of the system** We designed the CORUS system architecture using the AWS cloud service. When the user accesses the system, the web application, the API engine, the blockchain system, and the database communicate with each other in the server to exchange necessary data. In the process, we designed the server architecture in terms of the four aspects (reliability, availability, scalability and security).

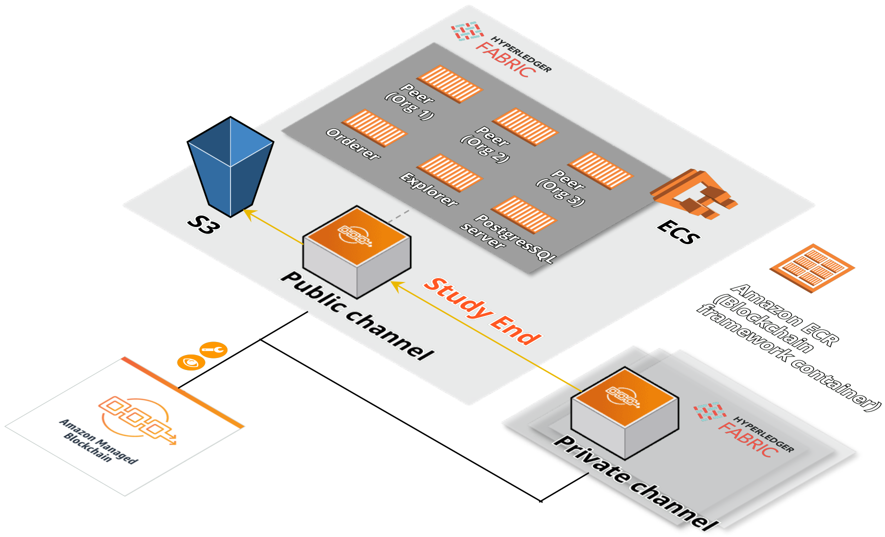


**Supplementary Figure 3. Conceptual diagram using the Amazon managed blockchain service** We configured the environment composed of a public channel and multiple private channels using the AWS services. The environment aimed to prevent data falsification.

**Supplementary Table 3. Features of the server architecture** We designed the architecture according to reliability, availability, scalability, and security, which demonstrates how well the server is designed.

| | |
|---|---|
| reliability | 1. Using two replicated availability zones with different server locations.<br>2. Distributed processing for each instance using load balancing.<br>3. Data transmission using message queues.[1] (SQS) |
| availability | 1. Reduce the difference in data transmission time due to distance by placing replicated servers in each region.[2] (CDN)<br>2. Maximize throughput with load balancing to minimize response time. |
| scalability | 1. Automatically adjust the number of instances by monitoring incoming traffic.[2,3] (Auto Scaling) |
| security | 1. Use https path based on secure shell layer(SSL) digital certificate, prevent A distributed denial-of-service(DDoS) attacks.[4] (Shield)<br>2. Separate private and public subnets, setting security rules for each instance.<br>3. Give the difference in instance access privileges within the administrators.[4,5] (IAM) |

### Technical specification

CORUS was developed as an open-source library, and the main language was developed using Java and JavaScript. Java used the Springboot framework to provide enterprise-class services to reliably handle numerous requests. And we developed to use rest application programming interface(API), Springboot's java persistence API(JPA) and GraphQL to service various requests called by Interface. The interface of the platform is composed of server side rendering(SSR) based on Node.js. The user interface(UI) is componentized using the Vue framework, and a responsive style is applied for mobile viewing. The database uses Postgresql, which is used to systematically store various data generated by the system. In addition, as mentioned in the text, we began development based on the predeveloped database of electronic data capture(EDC), OpenClinica's DB Schema, for systematic data storage and compatibility with other clinical specialized systems. Details of the languages and libraries used are listed in the supplementary table 4.

#### Blockchain to prevent data falsification

Preventing data falsification in vulnerable systems and the clinical research field often reports data falsifications. According to a National Institute of Health (NIH) study, 33% of researchers have experienced data falsification[6]. Blockchain is the most promising technology to prevent data distortion for any reason during the data storage. Characteristics of the blockchain in a clinical trial are immutable, traceable, and more trustworthy[7]. Recently, we demonstrated an integrated blockchain system to prevent data falsification for the crowdsourced data collection platform in participatory trials[8]. The system consists of a two-stage blockchain separated into a private chain for short-term data collection in progress and a public chain for long-term data storage from the completed study, respectively. The blockchain does not expose progressing data to participants to prevent the bandwagon effect and provide complete data to everyone to maintain transparency at the end of the study[9,10]. We used the blockchain system as the underlying data storage system of the participatory trial platform.

#### Data distribution

Data sharing can assist with ongoing research and positively impact derivative research[11,12]. In this regard, we expect that data sharing of the participants arouses a sustainable interest for ongoing or future studies. On the proposed platform, data sharing consists of two directions: transparently sharing research findings with the public and delivering them to a group of experts. Their detailed description is as follows. Sharing data with the public is aimed at transparently disseminating research findings and encouraging more participants to continue to access the platform based on trust. After the study is completed, anyone can view the study results through the platform interface. Users can easily share the results with other communities by using the SNS delivery function on the study results page. Furthermore, raw data can also be downloaded from the study. However, there is a problem with anonymity during this process[13] To solve this, it is now possible to use IDs that are randomly given when users participate in research[14]. Each table is stored separately in the database, and only relevant research information is provided to the researcher. For example, if a user with an ID of participates in a study number 2849, they will be assigned a random ID of 2849-1. Randomly assigned numbers are determined within the range of numbers of the target participants. Study creators can only view anonymous participant information. Participants can provide information on gender

**Supplementary Table 4.** **Service-specific language, library, and program details**

| | | | |
|---|---|---|---|
| API | Spring framewor | 2.0.4 | Apache License / Permissive license |
| | Graphql | 0.9.9 | MIT License / Permissive license |
| | Springfox | 2.9.2 | Apache License / Permissive license |
| | Lombok | 1.18.2 | MIT license / Permissive license |
| | Kotlin-logging | 1.4.9 | Apache License / Permissive license |
| | Commons-math | 3.6.1 | Apache License / Permissive license |
| | Commons-lang | 3.9 | Apache License / Permissive license |
| Interface | Node.js | 10.16.0 | MIT License / Permissive license |
| | Express | 4.15.2 | MIT License / Permissive license |
| | Vue | 2.6.10 | MIT License / Permissive license |
| | Jquery | 3.4.1 | MIT License / Permissive license |
| | Bootstrap | 4.3.1 | MIT License / Permissive license |
| | Datatables | 1.10.16 | MIT License / Permissive license |
| | Froala-editor | 2.8.4 | Commercial / Proprietary license |
| | Survey-creator | 1.1.0 | MIT License / Permissive license |
| | Survey-jquery | 1.1.0 | MIT License / Permissive license |
| | D3 | 5.12.0 | BSD licenses / Permissive license |
| | Chart.js | 2.7.2 | MIT License / Permissive license |
| | Chartist | 0.11.0 | MIT License / Permissive license |
| | Font-awesome | 4.7.0 | MIT License / Permissive license |
| | Mdi | 3.5.95 | None |
| | Flag-icon-css | 2.9.0 | MIT License / Permissive license |
| | I18next | 10.5.0 | MIT License / Permissive license |
| Database | Postgresql | 9.6.11 (AWS RDS) | PostgreSQL License / Permissive license |

and age in participatory studies, but this is optional, and even if such information is provided, identification of participants is very difficult in the absence of specific regional information[15]. Consequently, study results are transparent to the public, while personal information is opaque. This structure was developed based on Opensource EDC's Database Schema[16]. The proposed platform aims to collect reliable information. However, the data collected for detailed verification must be readily available to a group of experts. Convenient data utilization can only be achieved in a standardized format. Similarly, platforms such as PatientsLikeMe allow patients with rare diseases to share their treatment information, and clinical professionals can test the exact effect of shared information when interest in efficacy increases[17]. As such, in some cases, it is necessary to verify large-scale partial trial results using small-scale clinical trials precisely. This is a procedure that can express the effectiveness as efficacy[18]. This required a service that would independently exchange results or store and deliver clinical data. To this end, we have inherited the EDC function to receive data in the Operational Data Model (ODM) XML format that is compatible with the Clinical Data Interchange Standard Consortium[19,20].

#### System key features

CORUS consists of an API that provides data on demand and an interface that provides the desired screen through interaction with the user. The combination of these two provides a variety of features for the participant exam, and in addition to the core method described in the text, the step-by-step features that support it are shown in the supplementary figure 4.

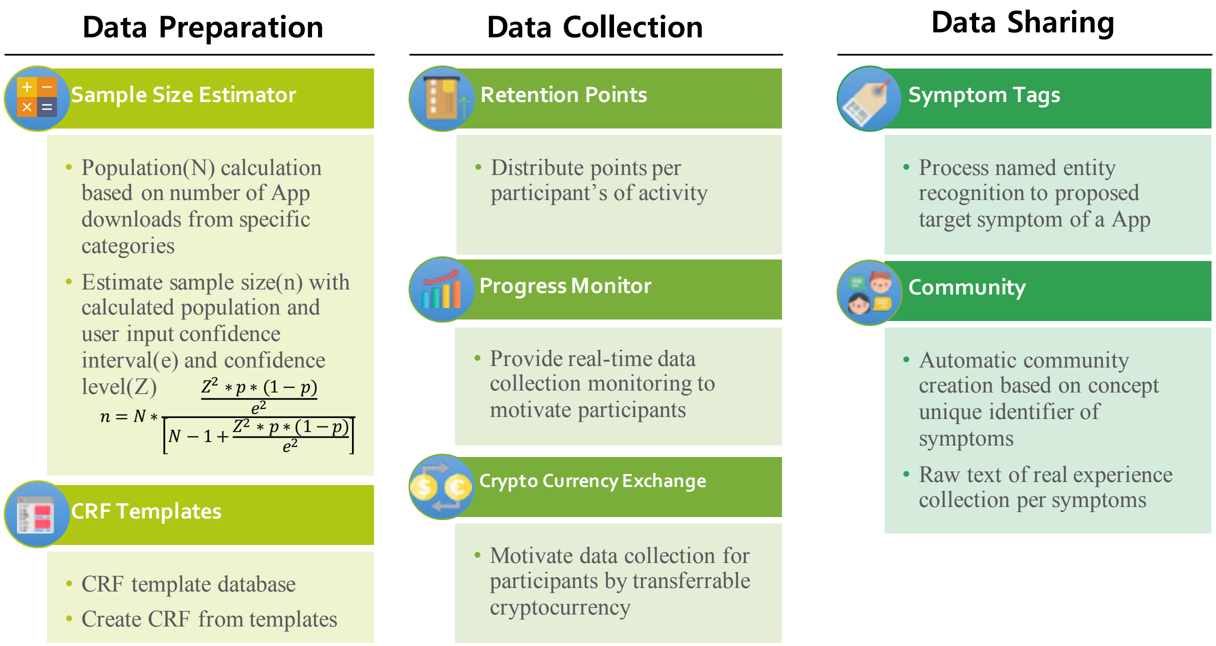


**Supplementary Figure 4. Tools to Facilitate the Collection of Data** CORUS offers a variety of features for data preparation, collection, and sharing. Modules for calculating sample size, eCRF templates shared by creators in predevelopedor advanced studies, make data preparation efficient. In addition to cryptocurrency, in the data storing phase, our platform provides activity points based on the user's activity, allowing the user to be interested in the system, access the system, and provide tools to monitor the progress of the research[21]. In addition, cryptocurrencies collected and distributed to individuals can be used on external exchanges to provide real value[22]. When a study is created, CORUS maps the words in Symptom to a biomedical concept to create a tag and uses that tag to create the DISQUS community[23]. This creates a place for many participants to discuss their questions actively.

#### System demo

Supplementary figure 5 demonstrates key features from study creation, conduct, and sharing and also describes changes in the blockchain that have been created by users within the platform. The platform version that is introduced in this video is 0.9 Beta, which might be slightly different from the current platform. If you want to check the latest screen, you should check https://corus.kaist.edu.

http://d2jjojlo5w7xa7.cloudfront.net/Supplementary_Video_1.mp4

**Supplementary Figure 5. CORUS demo** This video was released on June 11, 2019, at the AWS Public Sector Summit Washington, DC, session entitled "Blockchain for Science and Research," and is structured to showcase key system highlights.
